# Metabolite profiles across populations of Palmer amaranth (*Amaranthus palmeri*) highlight the specificity and inducibility of phytochemical response to glyphosate stress

**DOI:** 10.1101/2022.04.11.486891

**Authors:** Pawanjit Kaur Sandhu, Elizabeth Leonard, Vijay Nandula, Nishanth Tharayil

## Abstract

Modifications of the phytochemical profile form a vital component of physiological stress adaptation in plants. However, the specificity and uniqueness of phytochemical changes with respect to the identity of stressors is less known. Here, we investigated the commonality and specificity of metabolic perturbations induced by a specific stressor – glyphosate, and a general stressor – drought, across multiple glyphosate-resistant (GR) and -susceptible (GS) biotypes of a dominant agricultural weed, *Amaranthus palmeri*. In the absence of stress, the native metabolite profile of GS- and GR-biotypes was similar, and amplification of the EPSPS gene in GR-biotypes did not translate to a higher abundance of downstream metabolites. Further, glyphosate treatment initially inhibited the shikimate pathway in both GS- and GR-biotypes, from which the GR-biotypes recovered, indicating inducibility in the functionalization of the EPSPS enzyme. The accumulation of phenylpropanoids produced downstream of the shikimate pathway, was higher in GR-biotypes than GS-biotypes, with a preferential accumulation of compounds with higher antioxidant potential. However, this increase was not observed in response to drought treatment, where the metabolic perturbations were pervasive but limited in magnitude compared to glyphosate stress. Overall, while native phytochemistry of *A. palmeri* was similar irrespective of the level of glyphosate susceptibility, the specific stressor, glyphosate, imparted metabolic perturbations that were localized but higher in magnitude, while the specificity of phytochemical response to the general stressor, drought, was minimal. Taken together, these results suggest that, at the metabolic level, the glyphosate resistance mechanism in *A. palmeri* is partly induced and specific to herbicide stress.

**SIGNIFICANCE STATEMENT:** Understanding changes in physiology, especially those related to secondary metabolites with adaptogenic functions, is imperative to decipher the basis of stress adaptation in plants. This study provides critical information on native and stress-induced phytochemical differences between multiple glyphosate-resistant and -susceptible weed biotypes, thus, shedding light on the metabolome-level orchestration of gene amplification-mediated glyphosate resistance mechanism in an economically devastating weed, Palmer amaranth (*Amaranthus palmeri*).

## INTRODUCTION

Plants are adept at maneuvering the environment around them through changes in ontogeny, morphology, physiology, and biochemistry. A key underpinning of most physiological stress mitigation strategies in plants is the ability to produce a diverse array of phytochemicals, which facilitate plant adaptation through a multitude of functions, including serving as antioxidants, antimicrobials, signaling molecules, UV protectants, and feeding deterrents (Böttger et al., 2018). Perturbation of cellular metabolism in adverse growing conditions often alters the phytochemical composition of plants (Krasensky and Jonak, 2012), and some of these modifications enable the plants to circumvent stress. However, less known is the specificity of those chemical changes with respect to a stressor, i.e., whether the phytochemical changes in plants remain similar irrespective of the identity of the stressors. Further, it is unclear if conspecific plants (belonging to the same species) that differ in their stress mitigation capabilities would exhibit contrasting phytochemical responses to the same stressor i.e., if the phytochemistry could provide a cue to the differential ability of conspecifics to tolerate stress.

Herbicides, by disrupting biological processes, impart lethal stress to plants. However, natural variation in weed populations has facilitated the development of herbicide resistance in the biotypes of many weed species (Heap, 2021). The evolution of resistance in weeds to glyphosate, the most widely used herbicide in crop production systems, poses a major threat to global agriculture (Heap and Duke, 2018). One of the resistance mechanisms to this herbicide involves gene amplification of the 5-Enolpyruvylshikimate-3-phosphate synthase (EPSPS) enzyme, the target of glyphosate (Gaines et al., 2010). Glyphosate-resistant (GR) plants could possess more than 100-fold gene copies of EPSPS compared to glyphosate-susceptible (GS) plants (Gaines et al., 2010). EPSPS enzyme is a key driver of the shikimate pathway that results in the biosynthesis of aromatic amino acids (phenylalanine, tryptophan, and tyrosine), which in turn serve as precursors for a multitude of phytochemicals, including phenylpropanoids, alkaloids, hormones, and pigments (Maeda and Dudareva, 2012; Tohge et al., 2013). Since the regulation of the shikimate pathway is less evident in plants (Maeda and Dudareva, 2012; Zulet-González et al., 2020), the increased EPSPS gene copy numbers in GR plants could facilitate an increased abundance of downstream phytochemicals, many of which are known to protect plants from a variety of stressors (Deng and Lu, 2017; Sharma et al., 2019). Thus, the GR-biotypes with EPSPS gene amplification could produce a differential physiological and phytochemical response to stress compared to the GS-biotypes.

Based on the mode of action, environmental stressors can be broadly classified as those with a general or specific mode of action. Specific stressors disrupt a defined molecular target in plants (e.g., inhibition of specific enzymes), whereas general stressors perturb cellular homeostasis and plant physiology on a broader scale (e.g., abiotic stressors such as drought, heat, and cold that disrupt multiple enzymes and pathways) (Krasensky and Jonak, 2012). Consequently, the primary phytochemical response induced by a specific stressor is limited to the target pathway or pathways up/downstream of the target (e.g., inhibition of aromatic amino acid biosynthesis by glyphosate) (Amrhein et al., 1980; Maroli et al., 2015). On the other hand, the phytochemical response caused by general stressors reflects the perturbation of metabolic homeostasis across a multitude of pathways (Farooq et al., 2012; Krasensky and Jonak, 2012). Thus, depending on the specificity of response to stressors, the phytochemical response induced by glyphosate, a specific enzyme inhibitor, in GS- and GR-weed biotypes, could be different from the response induced by a general stressor such as drought that disrupts multiple physiological pathways in plants (Farooq et al., 2012).

Here, we tested these predictions in Palmer amaranth (*Amaranthus palmeri* [S. Wats].), a widespread GR weed species that exhibits EPSPS gene amplification as a mechanism of glyphosate resistance (Gaines et al., 2010). GR Palmer amaranth could possess up to 160 EPSPS gene copies that are known to yield higher transcript, protein, and enzyme activity levels even in the absence of glyphosate stress (Gaines et al., 2010; Ribeiro et al., 2014). However, the translation of the increased EPSPS enzyme activity to downstream aromatic amino acids and related metabolites is not known. Comparing the native- and herbicide-induced metabolism of GS- and GR-biotypes would not only increase our understanding of glyphosate resistance but could also facilitate the development of biomarkers that could predict glyphosate resistance in weed populations.

Metabolites, as the final products of genotype-environment interactions, are most closely related to phenotype (Patti et al., 2012). Thus, global metabolomic approaches that aim to characterize the complete set of small molecules in an organism present a promising avenue to investigate herbicide-induced physiological perturbations. Previous studies have utilized metabolomic approaches to capture the herbicide perturbations across different weed species (Maroli et al., 2015; 2016; 2018a; 2018b; Serra et al., 2015; Torres-García et al., 2018; Fernández-Escalada et al., 2019), including Palmer amaranth (Maroli et al., 2015, 2016; Fernández-Escalada et al., 2019). In these studies, each group (GS and GR) is represented by only one biotype. However, due to the outcrossing nature of Palmer amaranth (Chandi et al., 2012; Ward et al., 2013), the results obtained from the comparison of single GS- and GR-biotype might reflect genotypic variation between the two biotypes that could be unrelated to glyphosate resistance (Giacomini et al., 2014). Thus, to capture the true native and stress-induced metabolic differences between the GS- and GR-biotypes, genetic variation of dioecious Palmer amaranth needs to be considered (Giacomini et al., 2014; Küpper et al., 2018). Keeping this in view, we included multiple GS- and GR-biotypes of Palmer amaranth from diverse geographic locations across the US to investigate the metabolic perturbations caused by two different types of stressors, glyphosate (specific stressor) and drought (general stressor), using global metabolomic approaches.

Given the commonality of EPSPS amplification across GR-biotypes, we hypothesized that i) the populations of GR-biotypes would be distinguishable from those of GS-biotypes with respect to the native metabolite profile, ii) there will be commonalities in glyphosate-induced metabolic changes between the GS- and GR-biotypes, and when compared to their respective non-treated controls, the metabolic perturbations of GS-biotypes will be higher in magnitude than the GR-biotypes, and iii) compared to a general stressor (i.e. drought), the perturbations induced by a specific stressor (i.e. glyphosate) across biotypes, though higher in magnitude, will be limited to fewer metabolites due to the specific nature of herbicide inhibition.

## RESULTS

### Primary metabolites

Non-targeted analysis of polar metabolites using gas-chromatography – mass-spectrometry (GC-MS) resulted in the detection of 214 unique mass features. Of these, the putative identity of 54 mass features, including amino acids, organic acids, sugars, and related compounds, was established at confidence level 2 of the Metabolomics Standard Initiative (MSI) by matching the mass of the ion, fragmentation pattern, and retention index (RI) with the Kovats library [Table S1].

The sparse-Partial Least Square Discriminant Analysis (sPLS-DA) was used for dimensionality reduction of treatment effect on identified compounds. The first two components of sPLS-DA captured 40% of the variation in the data set. Axis-1 which explained 28% of the variation separated the glyphosate treatment from drought and control treatments across the biotypes, indicating a distinct primary metabolic profile of glyphosate treated GS- and GR-biotypes compared to control and drought treatments [Figure 1A]. This segregation was driven by a higher abundance of glucose-1-phosphate (G1P), shikimic acid, tyrosine, and tyramine in the glyphosate treatment, along with a lower abundance of tricarboxylic acid (TCA) cycle metabolites (oxalic acid, pyruvic acid, malic acid, succinic acid, and 2-oxoglutaric acid), sugars (fructose and glucose), organic acids (glyceric acid, threonic acid, and ascorbic acid), and amino acid alanine [Figure S1A]. The increased abundance of shikimic acid, one of the key drivers of this grouping, was related to an increase in the content of amino acids (tyrosine, glutamine, valine, and tryptophan), sugars and sugar alcohols (pentitol, G1P, tagatose, and myo-inositol) and ferulic acid and a decrease in TCA cycle metabolites (malic acid, citric acid, pyruvic acid, 2-oxoglutaric acid, succinic acid, and oxalic acid), organic acids (threonic acid, glyceric acid, 4-hydroxybutyric acid, and ascorbic acid), sugars (glucose and fructose), and amino acids (glutamic acid, alanine, and aspartic acid) [Fig S2]. The sPLS-DA axis-2 that explained 12% of the variation separated the glyphosate-treated GS- and GR-biotypes revealing that the primary metabolome of glyphosate-treated GS-biotypes was different from that of glyphosate-treated GR-biotypes [Figure 1A]. This clustering was based on the higher abundance of amino acids (tyramine, glycine, phenylalanine, and gamma-aminobutyric acid (GABA)), TCA cycle metabolites (fumaric acid, and cis-aconitic acid), and others (glycolic acid, dihydroxyacetone, 2-hydroxypyridine) in glyphosate-treated GR-biotypes, with a concomitant increase of shikimic acid, nicotinic acid, glutamine, myo-inositol, ribose, and glycerol in glyphosate-treated GS-biotypes [Figure S1B].

**Figure 1:**
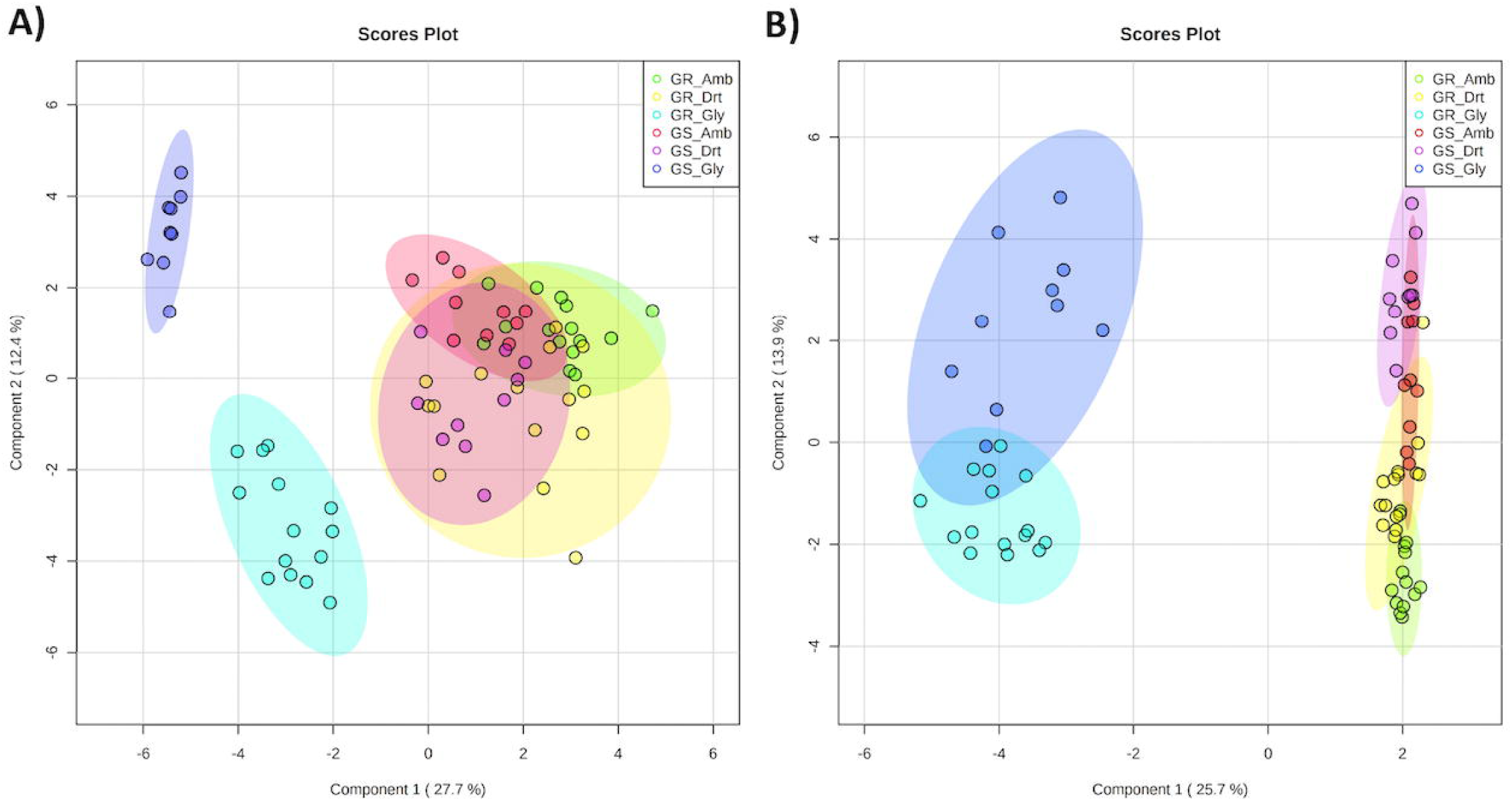
sPLS-DA score plot for the first two components of A) primary and B) secondary metabolomics data. The sparse-Partial Least Square Discriminant Analysis (sPLS-DA) score plot of the first two principal components of A). identified primary metabolites B.) top 2500 features from UHPLC-MS/MS analysis in glyphosate- (Gly), drought-(Drt) and control-(Amb) treatments of glyphosate-susceptible (GS)- and glyphosate-resistant (GR)- biotypes of Palmer amaranth. Within each group (GS & GR) the biotype identity is a random factor. The ellipses represent a 95% confidence interval.

Compared to glyphosate treatment, drought treatment resulted in a lower degree of variation in the metabolic pool across the biotypes. Similar to glyphosate treatment, as evident from the hierarchical clustering analysis [Figure 2], the drought and control treatment had an overriding effect on the metabolite composition over the identity of the biotypes. The drought treatment resulted in an increased abundance of sugars (glucose, galactose, and sucrose), amino acids (valine, tryptophan, isoleucine, and GABA), organic acids (glycolic acid, threonic acid, 4-hydroxybutyric acid, and oxalic acid), and other metabolites (dihydroxyacetone, N-acetyl hexosamine, 2-hydroxypyridine) while the abundance of amino acids (aspartic acid, glutamic acid, alanine), organic acids (ascorbic acid, glyceric acid), TCA cycle metabolites (malic acid, succinic acid, 2-oxoglutaric acid, pyruvic acid), and fructose was reduced compared to non-treated control treatment [Figure 2].

**Figure 2:**
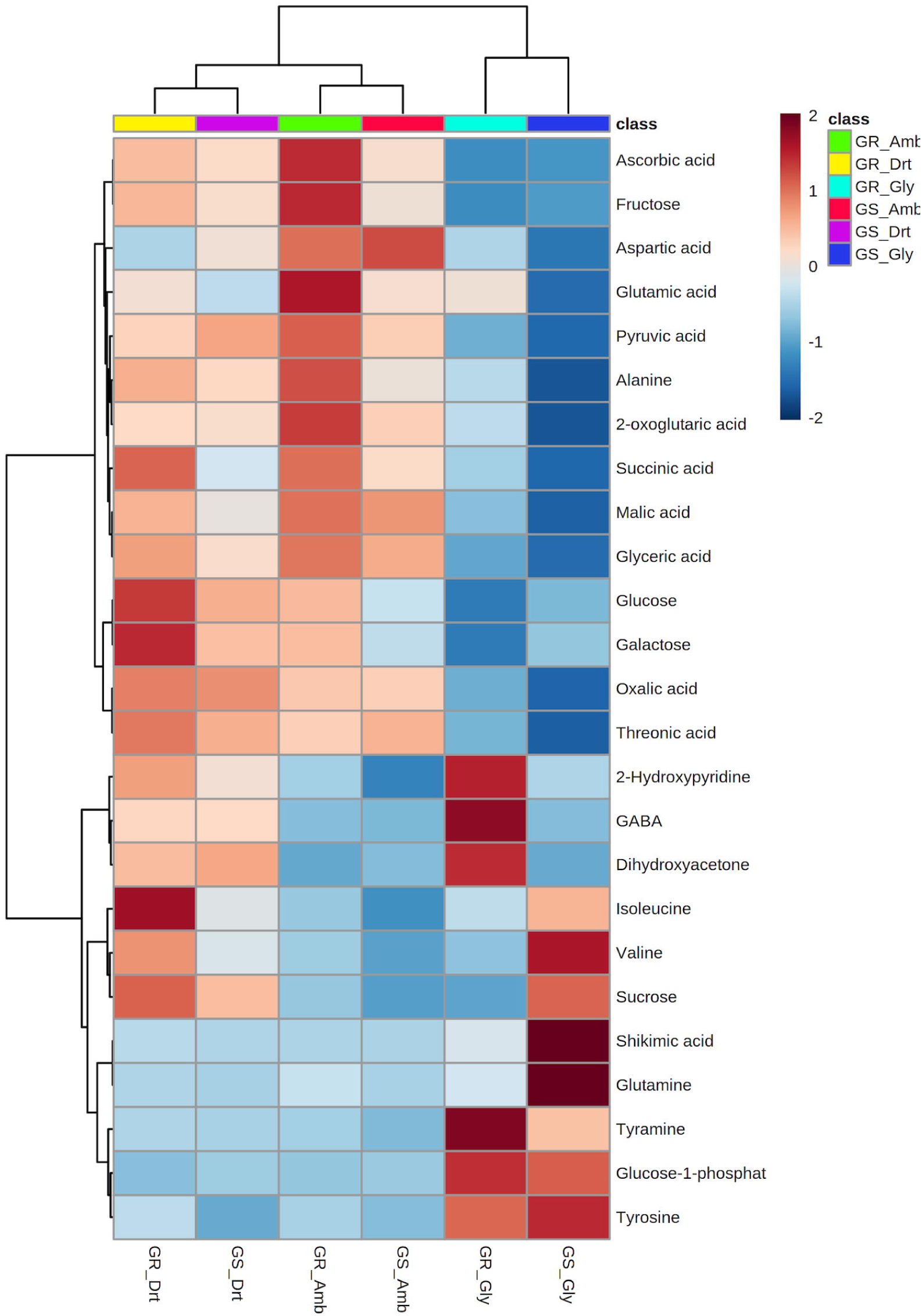
Heatmap and two-way hierarchical clustering of identified primary metabolites. Heatmap and two-way hierarchical clustering of identified primary metabolites in glyphosate- (Gly), drought- (Drt) and control- (Amb) treatments of glyphosate-susceptible (GS)- and glyphosate-resistant (GR)- biotypes of Palmer amaranth. Within each group (GS & GR) the biotype identity is a random factor.

#### a. Innate differences between the primary metabolome of GS- and GR-biotypes

Based on the primary metabolic profile, non-treated GS-biotypes clustered distinctly from that of GR-biotypes [Figure S3]. This clustering was driven by the higher abundance of amino acids (glutamic acid, 5-oxoproline, alanine, serine, glutamine, isoleucine, and glycine), sugars (glucose, methyl glucopyranoside, galactose, and fructose), and organic acids (ascorbic acid and 2-oxoglutaric acid) in the GR-biotypes and N-acetylhexosamine and dihydrouracil in the GS-biotypes [Figure S3]. These differences were also significant in univariate analysis, although the log_2_ fold change was less than the chosen threshold of one [Figure S4, Table S2].

#### b. Differential effect of glyphosate treatment on the primary metabolome of GS- and GR-biotypes

Glyphosate treatment perturbed the primary metabolism (TCA cycle, amino acids, and sugars) in both GS- and GR-biotypes, although the magnitude of disruption was biotype-dependent. Both GS- and GR-biotypes showed a decrease in the abundance of all TCA cycle metabolites except fumaric acid and cis-aconitic acid after exposure to glyphosate [Figure 3, Table S2]. However, the reduction of these metabolites was higher for GS-biotypes than GR-biotypes, indicating a stronger effect of glyphosate on GS-biotypes. Further, log_2_ fold change in the GR-biotypes for citric acid, succinic acid, and 2-oxoglutraic acid was lower than the chosen threshold value of one. In addition, GR-biotypes showed an accumulation of cis-aconitic acid and depletion of total sugar metabolites [Figure S5] which was not observed in GS-biotypes.

**Figure 3:**
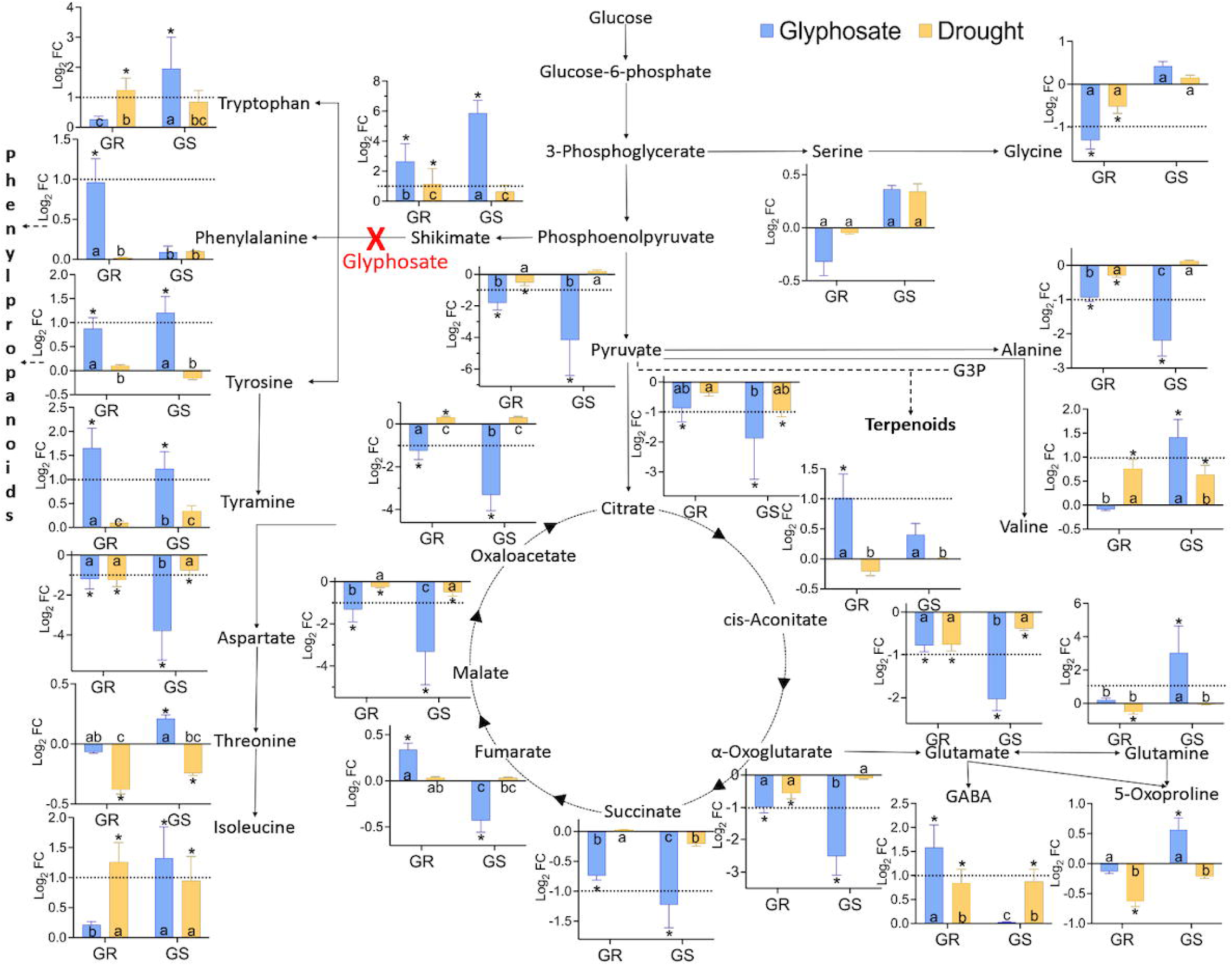
Effect of glyphosate and drought treatment on TCA cycle metabolites and amino acids. Log_2_ fold change in tricarboxylic acid (TCA) cycle **metabolites** and amino acids in glyphosate- and drought-treated glyphosate-susceptible (GS)- and glyphosate-resistant (GR)- biotypes of Palmer amaranth. The red cross shows the point of inhibition of glyphosate. The bar represents mean ± SD. Asterisk denotes the significant difference of treatment compared to control at p < 0.05 and FDR <0.05 (pairwise comparisons). The dotted line in the graphs indicates the log_2_ fold change threshold of |1|, which was chosen as significant. Bars with different letters are significantly different from each other. Within each group (GS & GR) the biotype identity is a random factor.

Similar to TCA cycle metabolites, amino acids were perturbed more in glyphosate-treated GS-biotypes than GR-biotypes. In GR-biotypes, glyphosate treatment resulted in a reduction of non-aromatic amino acids (aspartic acid and glycine) along with an increase in GABA and aromatic amino acid tyramine. In GS-biotypes, fold change in the abundance of aspartic acid was higher while glutamic acid, alanine, valine, isoleucine, glutamine (non-aromatic amino acids), tyrosine, and tryptophan (aromatic amino acids) were also accumulated [Figure 3].

Shikimic acid accumulation is a metabolic marker associated with glyphosate-induced inhibition of the EPSPS enzyme. Accordingly, shikimic acid was higher in glyphosate treatment compared to drought and control treatments in both biotypes, and the accumulation of shikimic acid was significantly higher in GS-biotypes than GR-biotypes [Figure 3, Table S2].

#### c. Differential effect of drought treatment on the primary metabolome of GS- and GR-biotypes

The impact of drought treatment on the primary metabolism of GS- and GR-biotypes was minimal compared to glyphosate treatment. None of the TCA cycle metabolites were affected across the biotypes at the chosen threshold of one. However, there was a slight reduction (log_2_ fold change < 1) in the abundance of malic acid and citric acid in the GS-biotypes and oxalic acid, malic acid, 2-oxoglutaric acid, and pyruvic acid in the GR-biotypes [Figure 3]. Further, the impact of drought treatment on amino acids was observed only in GR-biotypes with an accumulation of isoleucine and tryptophan and reduction of aspartic acid [Figure 3] while sugars were perturbed specifically in GS-biotypes (accumulation of 1-methylgalactose and depletion of fructose) [Figure S5]. Drought-treated GR-biotypes also showed an increase in shikimic acid content, albeit to a much lower extent than glyphosate treatment – a two-fold increase in drought treatment compared to six-fold accumulation in the glyphosate treatment [Figure 3, Table S2].

### Secondary metabolites

To understand the effect of glyphosate resistance and stress treatments on secondary metabolome, sPLS-DA dimension reduction was performed on top 2,500 unique mass features obtained from ultra-high pressure liquid-chromatography – tandem mass-spectrometry (UHPLC-MS/MS) analysis. The first two components of the sPLS-DA explained 40% of the variation in the dataset [Figure 1B]. Similar to the primary metabolome, the component axis-1 explaining 25.7% variation separated the glyphosate treatment from drought and control treatments across the biotypes. The component axis-2 that explained 13.9% of the variation differentiated GS- and GR-biotypes within the glyphosate treatment. This indicates a distinct secondary metabolic profile of glyphosate-treated GS- and GR-biotypes from respective control and drought treatments as well as from each other. On the other hand, the drought treatment clustered with control treatment in both GS- and GR-groups [Figure 1B].

To date, there are no published reports of secondary metabolite identification in Palmer amaranth. In this study, we manually curated the identities of 162 secondary metabolites (including phenylpropanoids and terpenoids) at confidence-level 2 and 3 of MSI by cross comparing the accurate mass (<5ppm) and fragmentation spectra of compounds with that of online databases (Mass bank of North America (MoNA), mzCloud, Flavanoid Search, Human Metabolome Database (HMDB)) and literature search (Akimoto et al., 2017; Li et al., 2018; Ma et al., 2019; Tsugawa et al., 2019). The details of all the identified metabolites are provided in a supplementary table [Table S3].

### Phenylpropanoids and Phenolic compounds

124 metabolites including flavonoids, cinnamic acid derivatives, coumarins, and phenolic compounds were identified based on their characteristic molecular fragment pattern and neutral losses [Table S3] (Akimoto et al., 2017; Li et al., 2018; Fernández-Poyatos et al., 2019; Ma et al., 2019; Tsugawa et al., 2019; Manyelo et al., 2020).

Identification of hydroxycinnamic acid amides was based on the annotation method proposed by Li et al. (2018). Further, cinnamic acid derivatives and flavonoids accounted for more than 90% of the identified phenylpropanoids [Figure 4]. Based on diversity (in terms of the number of compounds within each class), cinnamic acid derivatives (66.1%) were the most abundant class, followed by flavonoids (25.8%) and coumarins (4%). However, based on the abundance (in terms of peak intensities), the flavonoids (61.6%) were the highest abundance class, followed by cinnamic acid derivatives (35.9%) [Figure 4]. Thus, Palmer amaranth exhibited a higher diversity of cinnamic acid derivatives, but a higher content of flavonoids in this study. Flavonoid profile was mainly dominated by glycosides of flavonols (quercetin (41.9%) and kaempferol (16.3%)) with a low abundance of other flavonoids (naringenin chalcone, naringenin, myricetin, isorhamnetin, and dihydroquercetin). Within cinnamic acid derivatives, ferulic acid derivatives (20.2%) were the most abundant subclass, followed by coumaric acid derivatives (8.6%) and caffeic acid derivatives (4.5%). Other identified phenylpropanoids included coumarin glycosides (5), phenolic glycosides (4), and syringic acid. Amaranthaceae-specific betalains were not observed.

**Figure 4:**
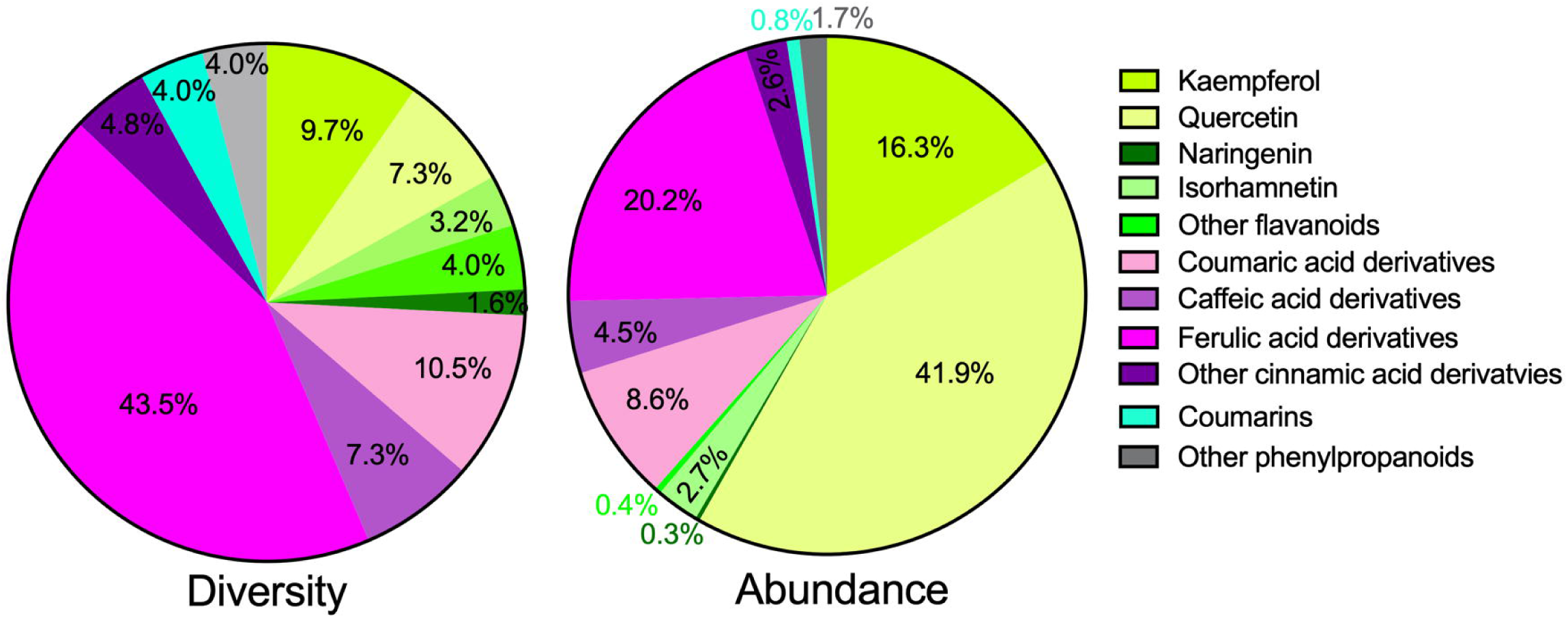
Composition of identified phenylpropanoids and phenolic compounds based on diversity and abundance of various classes. The values indicate the percentage of a particular class based on diversity and abundance. The classes within cinnamic acid derivatives are represented by shades of green color while those within flavonoids are represented by shades of pink color. Diversity was calculated as the number of compounds identified in a class while summed intensity was used as a measure of abundance.

### Terpenoids

Thirty-two terpenoid glycosides were identified based on accurate mass and characteristic fragmentation patterns (Fiorentino et al., 2006; Nedialkov et al., 2012; Ma et al., 2019) [Table S3]. All but three of the identified terpenoids were triterpenoid glycosides. The most common glycosylation moiety was hexuronic acid based on a neutral loss of 176.032 amu, followed by pentoside (132.043 amu) and hexoside (162.053 amu) [Table S7]. The identified triterpenoid aglycones were oleanolic acid, hederagenin, arjunolic acid, akebonoic acid, normedicagenic acid, medicagenic acid, and dihydroxydioxooleanoic acid [Table S7]. These triterpenoid aglycones have been identified previously across various species in the Amaranthaceae family (Mroczek, 2015). The position of glycosylation at the triterpenoid ring was determined based on the differential fragmentation pattern of C-3 vs. C-28 glycosylated triterpenoids (Ma et al., 2019). Other identified terpenoid classes were: sesquiterpenoid glycosides (2) and a tetraterpenoid glycoside (1).

In addition to phenylpropanoids and terpenoids, four nitrogenous glycosides (vicine, hypoxanthine pentoside, indoxyl hexopyranoside_427.148_9.13, and indole glycoside_367.127_10.02) and two jasmonic acid-derived glycosides (tuberonic acid hexoside, and isocucurbic acid hexoside) were also identified (Rizzello et al., 2016; Tzachristas et al., 2021) [Table S3].

### CANOPUS classification

Since two-thirds of the captured UHPLC-MS/MS profile could not be annotated, a machine learning-based software, CANOPUS (Dührkop et al., 2021), was employed to predict the chemical classification of the mass spectra. Prior to analysis with CANOPUS, mass spectra were filtered to remove insource fragments, adducts, and peaks with low abundance producing an abundance matrix of 3571 unique mass features. Of the filtered features, 75% were classified by CANOPUS into thirteen superclasses [Figure 5], all belonging to kingdom organic compounds [Table S5]. Five superclasses, including lipids and lipid-like molecules (31.3%), organic acids and derivatives (25.4%), organic oxygen compounds (16.2%), phenylpropanoids and polyketides (10.9%), and benzenoids (7.8%), accounted for more than 90% of compounds [Figure 5]. The superclasses were further categorized up to level 5 of ClassyFire (Djoumbou Feunang et al., 2016), including classes, subclasses, and level 5, respectively [Table S5]. Prenol lipids were the highest abundant group (22.5%) of secondary metabolites, which was mainly dominated by triterpenoid glycosides and triterpenoids subclasses consistent with the composition observed for identified terpenoids. Similarly, within phenylpropanoid and polyketides, cinnamic acid derivatives were the most abundant class, followed by flavonoids and coumarins and derivatives [Figure 5].

**Figure 5.**
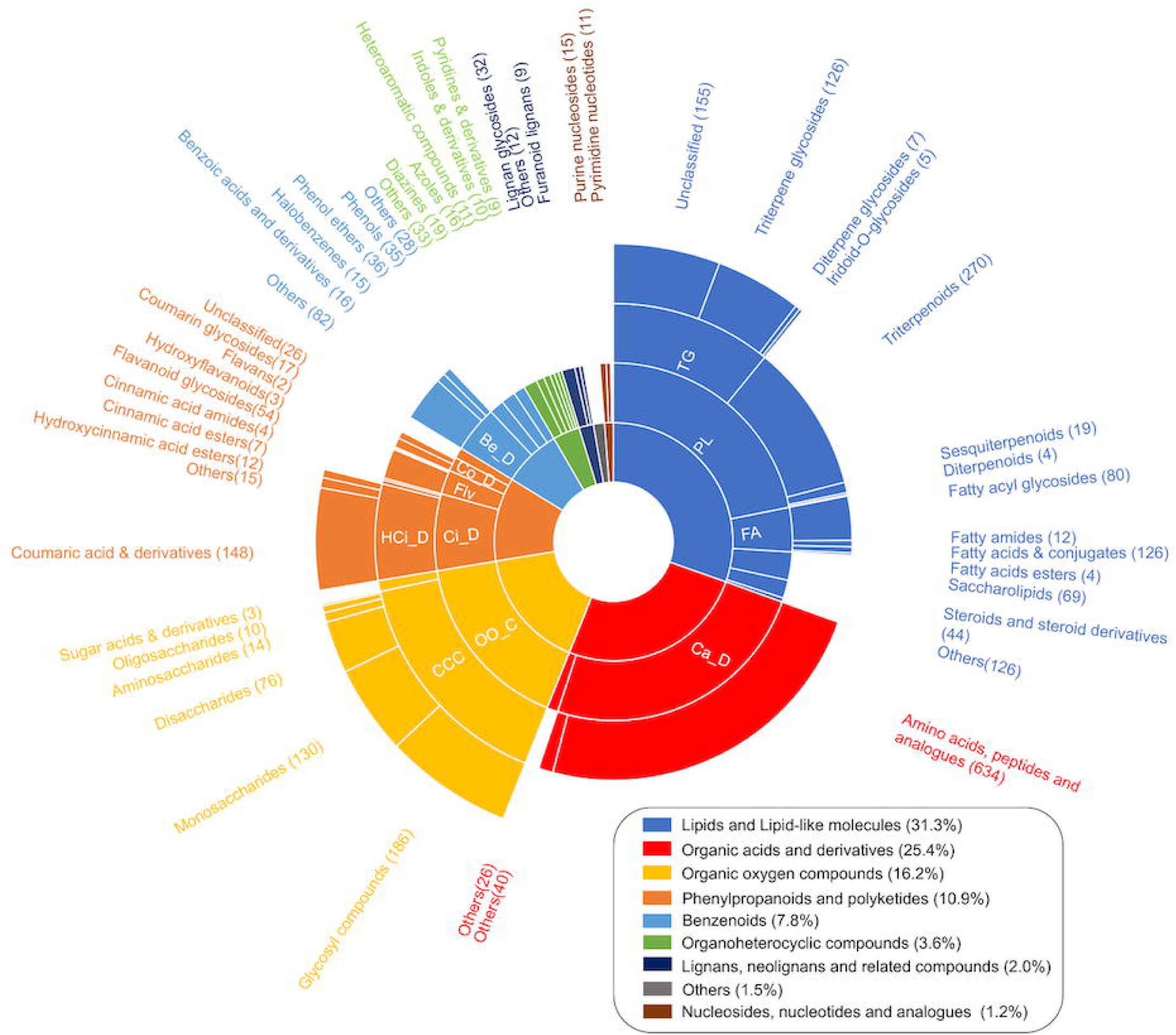
CANOPUS classification of the filtered features from UHPLC-MS/MS analysis. The numbers in the brackets of a subclass in the sunburst plot represent the number of compounds belonging to the respective subclass while the percentage in the brackets of superclass in legend represents the percentage of compounds belonging to respective superclass. Abbreviations: Be_D: Benzene and substituted derivatives, Ca_D: Carboxylic acids and derivatives, CCC: Carbohydrates and carbohydrate conjugates, Ci_D: Cinnamic acids and derivatives, Co_D: Coumarins and derivatives, FA: Fatty acyls, Flv: Flavanoids, HCi_D: Hydroxycinnamic acids and derivatives, OO_C: Organoooxygen compounds, PL: Prenol lipids, TG: Terpene glycosides.

#### a. Innate differences between the secondary metabolome of GS- and GR-biotypes

The abundance of more than 80% of the metabolites from CANOPUS analysis was similar in the non-stressed GS- and GR-biotypes. Further, contrary to our hypothesis, GS-biotypes had twice the number of phytochemicals with higher native abundance compared to the GR-biotypes in all the CANOPUS superclasses, including lipids and lipid-like molecules, organic acids and derivatives, organic oxygen compounds, phenylpropanoids and polyketides, benzenoids, as well as unclassified compounds [Figure 6A]. The only exception was lignans, neolignans, and related compounds superclass, where seven compounds had a higher abundance in the non-treated GR-biotypes compared to one in the non-treated GS-biotypes [Figure 6A].

**Figure 6.**
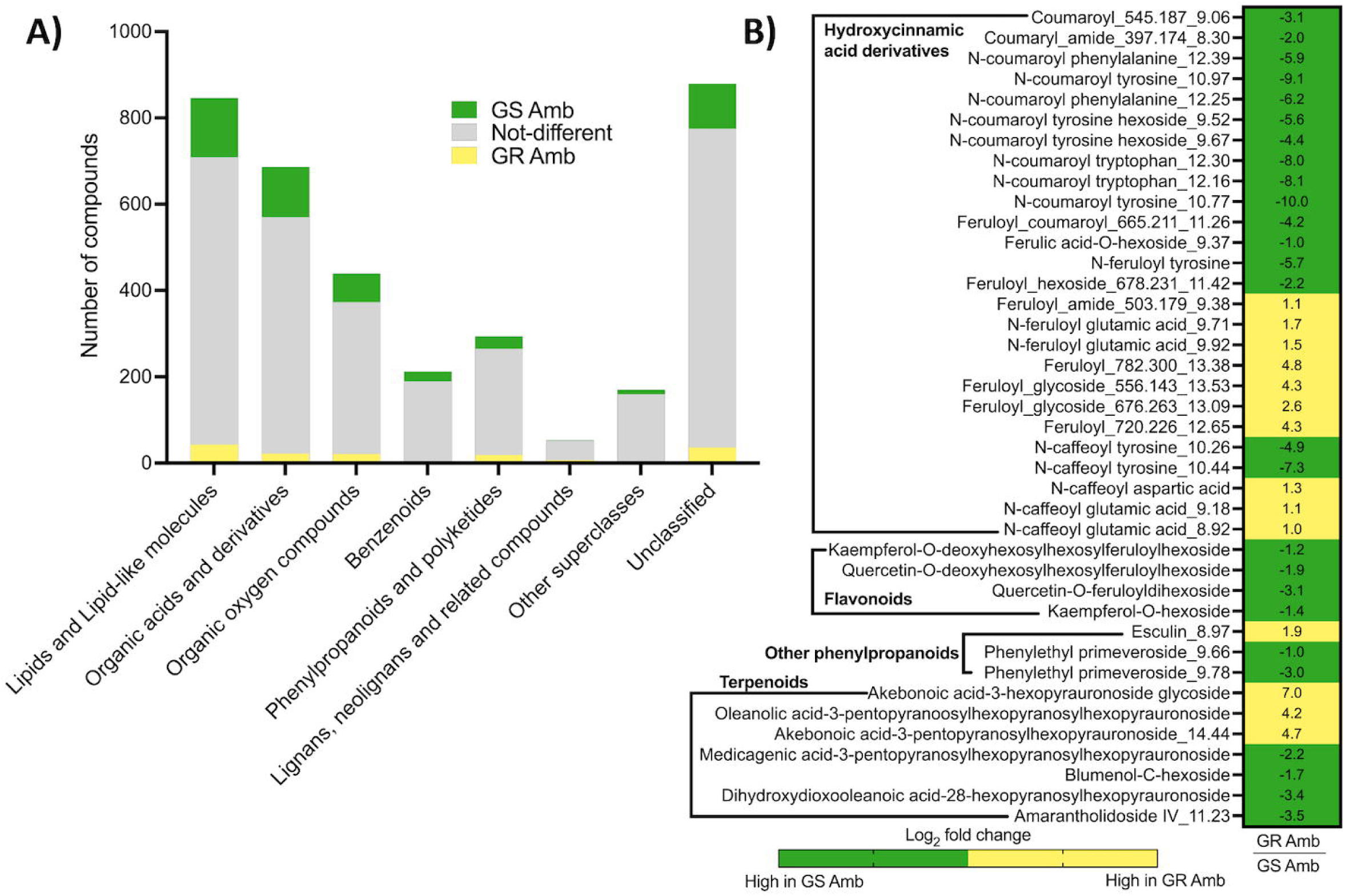
Native differences between glyphosate-susceptible (GS-) and glyphosate-resistant (GR-) biotypes. Compositional differences between the GS- **and** GR-biotypes of Palmer amaranth in the control (Amb) treatment. A). At the CANOPUS superclass level: The green and yellow shaded area in the bar respectively represents the number of compounds higher in the control treatment of GS (GS Amb) and GR (GR Amb) biotypes (Log_2_ fold change > |1|, p <0.05, FDR <0.05). The gray shaded area represents metabolites that were not different between the two treatments. B) Log_2_ fold change (GR Amb/GS Amb) plot of identified secondary metabolites that were significantly different (Log_2_ fold change > |1|, p <0.05, FDR <0.05) between the GS Amb and GR Amb treatments. Values in each cell are the GR Amb/GS Amb log_2_ FC values. Within each group (GS & GR) the biotype identity is a random factor.

Consistent with CANOPUS analysis, the abundance of more than 75% of the identified secondary metabolites was not different between the control GS- and GR-biotypes, and GS-biotypes had twice the number of phenylpropanoids (22) with higher native abundance compared to the GR-biotypes (11). Specifically, non-treated GS-biotypes were high in aromatic amino acid conjugates of coumaric acid, caffeic acid, and ferulic acid, other ferulic acid derivatives, glycosides of kaempferol and quercetin, as well as other phenolic glycosides [Figure 6B]. The GR-biotypes showed a higher abundance of glutamic acid conjugates of ferulic acid and caffeic acid, N-caffeoyl aspartic acid, other ferulic acid derivatives, and coumarin esculin_8.97 [Figure 6B]. The proportional difference (22 vs. 11) observed for phenylpropanoids was not observed for identified terpenoids – GS-biotypes showed an innately higher abundance of four terpenoids (including a sesquiterpenoid, a tetraterpenoid, and two triterpenoid glycosides) compared to three triterpenoid glycosides in the GR-biotypes [Figure 6B, Table S4].

#### b. Differential effect of glyphosate treatment on the secondary metabolome of GS- and GR-biotypes

In GR-biotypes, more than 1100 metabolites (33% of the total) increased in response to glyphosate compared to 900 metabolites (25% of the total) in the GS-biotypes [Figure 7A]. Major contributors to this difference were phenylpropanoids and polyketides and organic oxygen compounds, where GR-biotypes had four times the metabolites that accumulated in GR-biotypes but were unperturbed in GS-biotypes [Figure 7A]. Further, on average, 80% of the metabolites that accumulated in GS-biotypes also increased in the GR-biotypes. Contrastingly, only 5% of the captured metabolites decreased in response to glyphosate, indicating that the glyphosate-induced phytochemical response was driven by accumulating metabolites across the biotypes. Interestingly, the downregulated metabolic profile of glyphosate-treated GS- and GR-biotypes was deficient in the phenylpropanoids and polyketides and organic oxygen compounds superclasses which had a higher abundance of upregulated metabolites [Figure 7A].

**Figure 7.**
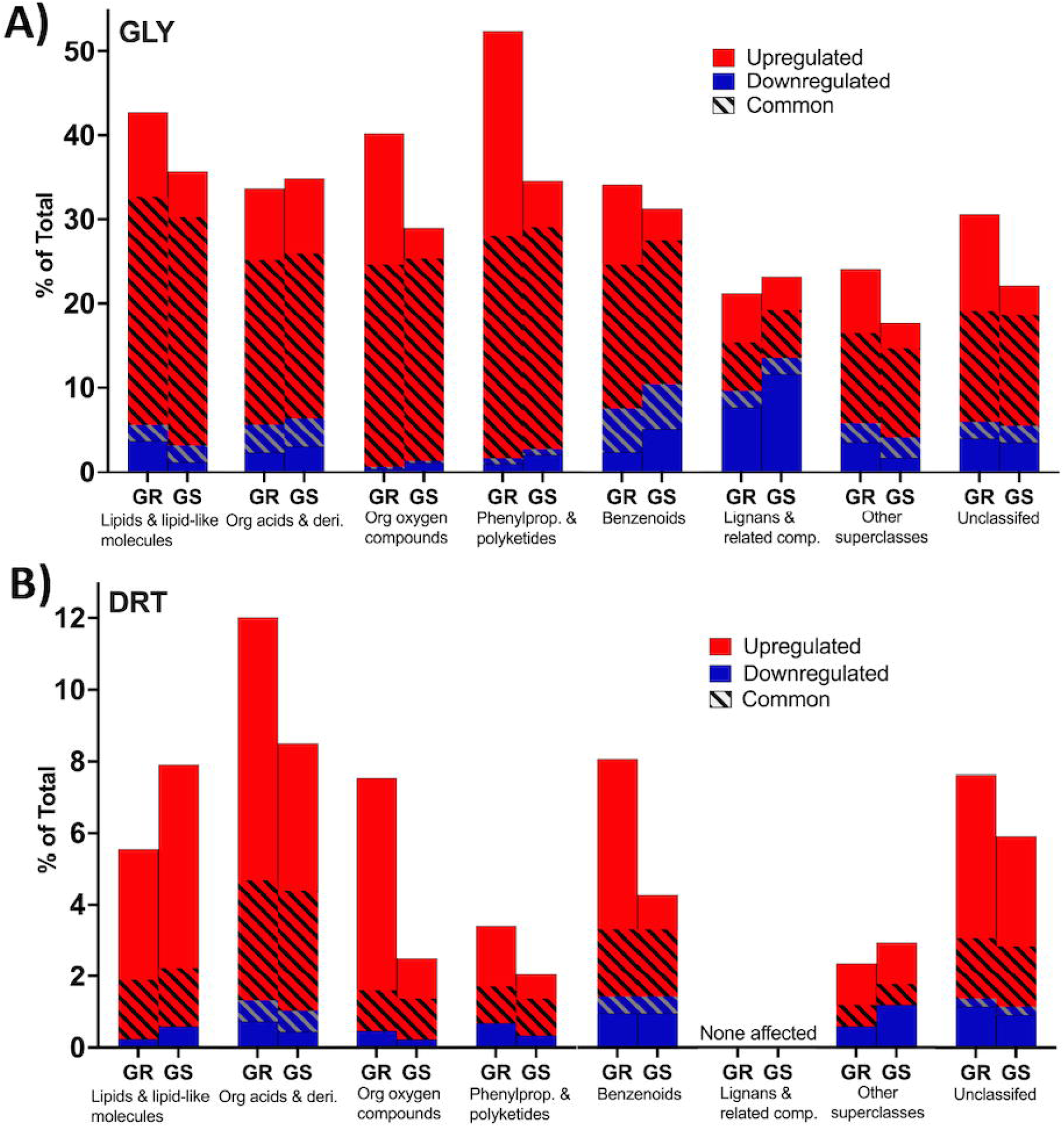
Effect of A) glyphosate and B) drought treatment on UHPLC-MS/MS analyzed profile. Bar plots displaying the effect of A) Glyphosate B) Drought on the UHPLC-MS/MS analyzed profile of GS- and GR-biotypes of Palmer amaranth in the superclasses predicted by CANOPUS classification. The X-axis represents various superclasses and the Y-axis represents the percentage of compounds up(red)- or down(blue)-regulated (p<0.05, FDR <0.05, log_2_FC > |1|) in treated (Glyphosate or Drought) glyphosate-susceptible (GS-) and glyphosate-resistant (GR-) biotypes with respect to controls. The shaded area within each bar represents the compounds that were affected in both groups. Within each group (GS & GR) the biotype identity is a random factor.

The trends observed from CANOPUS analysis were also true for identified secondary metabolites: glyphosate treatment resulted in the accumulation of a greater number of phenylpropanoids in GR-biotypes (79) than GS-biotypes (55), and more than 85% of the phenylpropanoids that accumulated in glyphosate-treated GS-biotypes also increased in the GR-biotypes [Figure 8, Table S8]. The phenylpropanoids that increased across the groups were comprised of derivatives of ferulic acid, caffeic acid-O-hexoside, and glycosides of quercetin, kaempferol, isorhamnetin, dihydroquercetin, and scopoletin [Figure 8]. GR-biotypes also reported an accumulation of additional ferulic acid derivatives that was not observed in the GS-biotypes. In fact, of the 54 identified ferulic acid derivatives, 49 compounds accumulated in the glyphosate-treated GR-biotypes [Figure 8]. Further, coumaric acid derivatives, glycosides of quercetin, isorhamnetin, and myricetin also increased specifically in the glyphosate-treated GR-biotypes. The phenylpropanoids that accumulated only in GS-biotypes included derivatives of coumaric acid (1), ferulic acid (1), and glycosides of kaempferol (3) and naringenin (2). GS-biotypes also showed a reduction in the abundance of coumaroyl phenylalanine conjugates, naringenin chalcone, and two other hydroxycinnamic acid derivatives, which was not observed in the GR-biotypes [Figure 8, Table S4].

**Figure 8:**
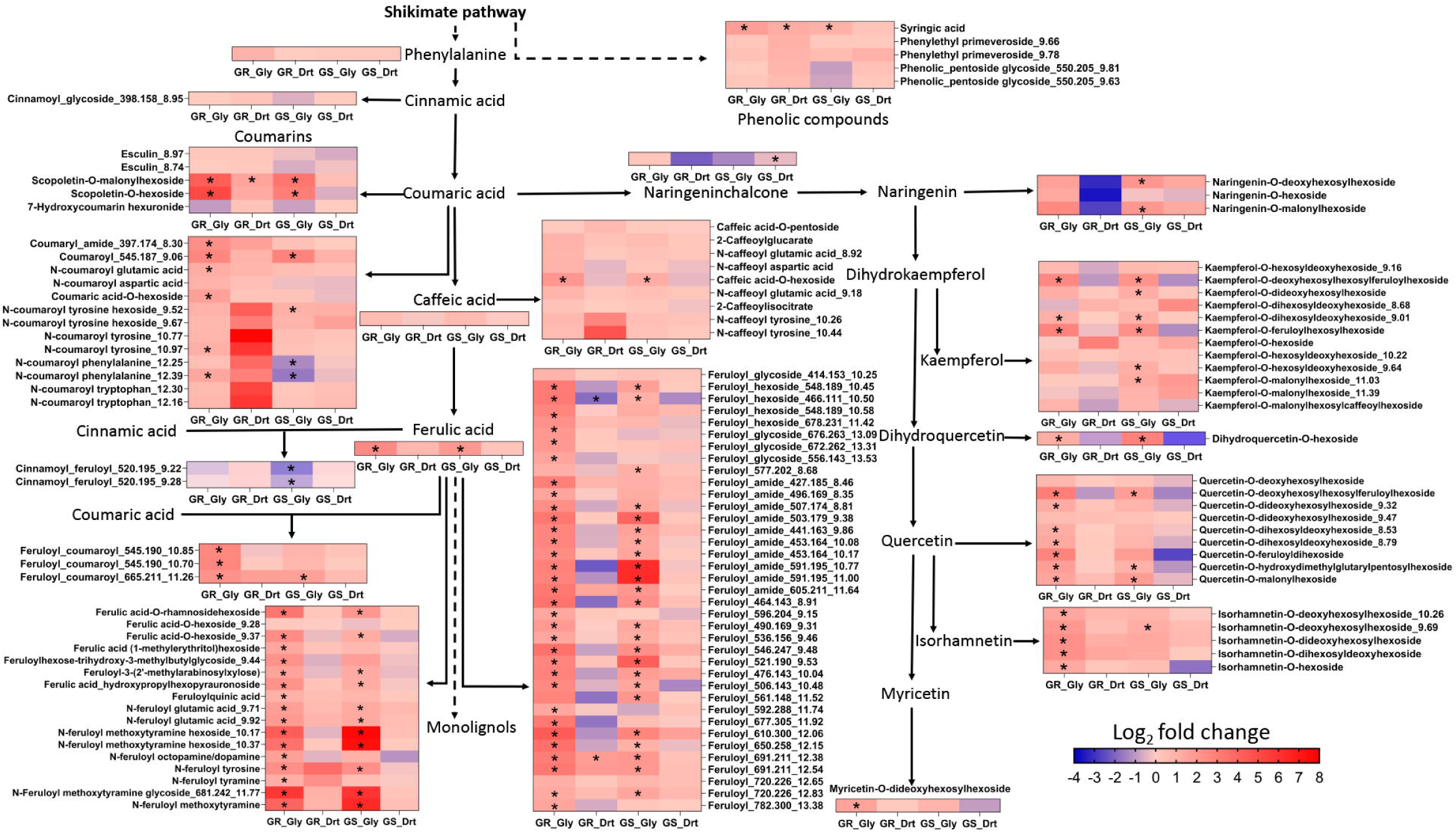
Effect of glyphosate and drought treatment on identified phenylpropanoids and phenolic compounds. Log_2_ fold change of identified phenylpropanoids and phenolic compounds in glyphosate- (Gly) and drought- (Drt) treated glyphosate-susceptible (GS)- and glyphosate-resistant (GR)- biotypes of Palmer amaranth compared to respective controls. The significantly affected metabolites (p<0.05, Log_2_ fold change ≥ |1|, FDR <0.05) are denoted with an asterisk. Non-identified compounds are annotated with the accurate mass followed by retention time. Within each group (GS & GR) the biotype identity is a random factor.

In contrast to the phenylpropanoids, the effect of glyphosate treatment on the abundance of terpenoids was similar across the groups [Figure 9]. Twenty terpenoids accumulated in response to glyphosate, of which seventeen were common between GS- and GR-biotypes, including sixteen triterpenoid glycosides and a sesquiterpenoid [Figure 9, Table S4]. The other three triterpenoid glycosides accumulated in a group-specific manner in GS- and GR-biotypes, while none of the terpenoids showed depletion across the biotypes.

**Figure 9:**
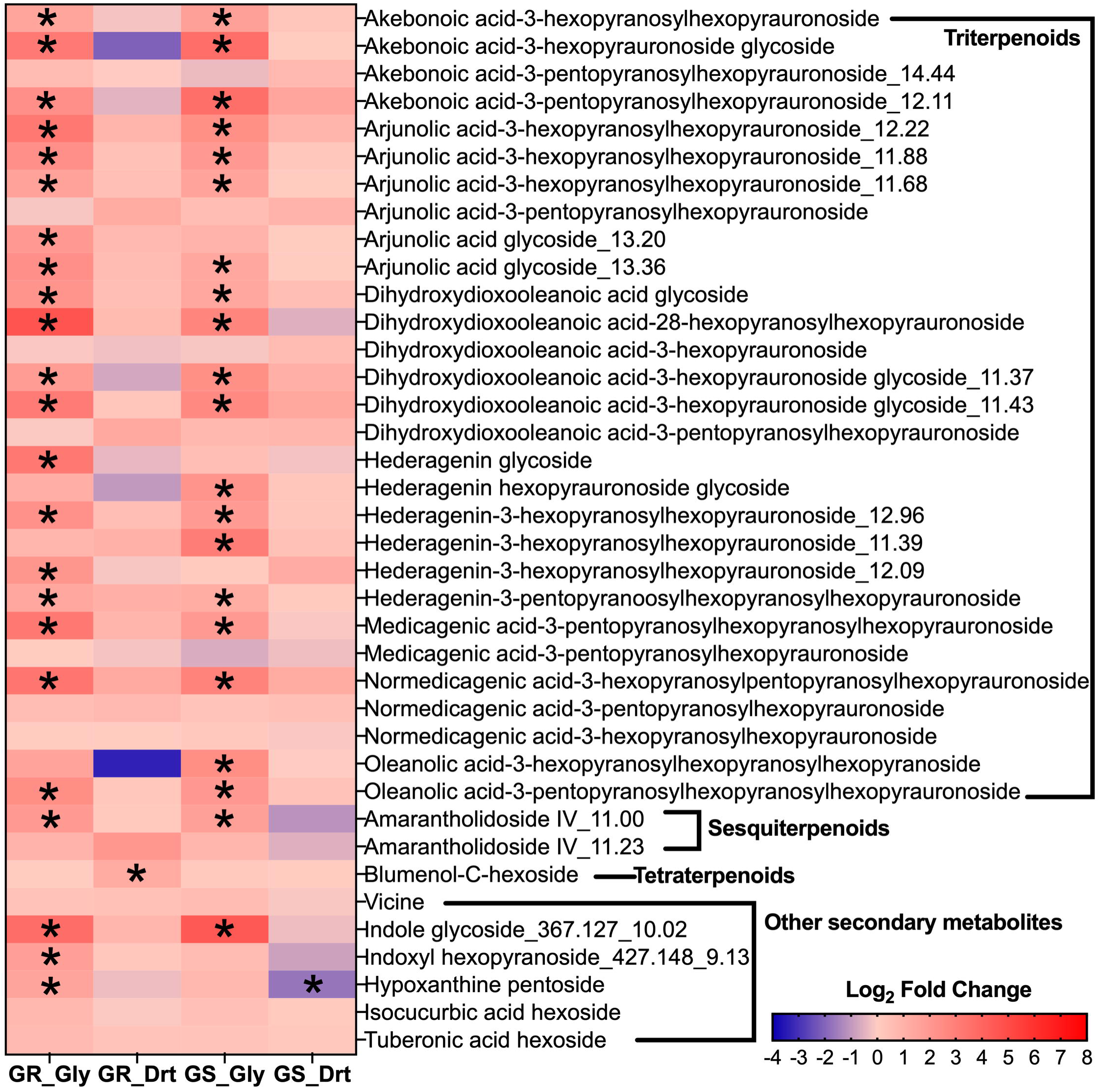
Effect of glyphosate and drought treatment on identified terpenoids and other secondary metabolites. Heatmap showing log_2_ fold change of identified terpenoids and other secondary metabolites in glyphosate- (Gly) and drought- (Drt) treated glyphosate-susceptible (GS-) and glyphosate-resistant (GR-) of Palmer amaranth compared to respective controls. The significantly affected metabolites (p<0.05, Log_2_ fold change ≥ |1|, FDR <0.05) are denoted with an asterisk. Within each group (GS & GR) the biotype identity is a random factor.

Both the GR- and GS-biotypes also showed a glyphosate-induced accumulation of indole hexoside_367.127_10.02, while indoxyl hexopyranoside_427.148_9.13 and hypoxanthine pentoside increased only in the GR-biotypes [Figure 9].

#### c. Differential effect of drought treatment on the secondary metabolome of GS- and GR-biotypes

Similar to primary metabolism, the effect of drought treatment on the secondary metabolome of Palmer amaranth was limited [Figure 7B, 8, 9]. Within CANOPUS analysis, only 6 and 7% of the metabolites were disrupted by drought treatment in GS- and GR-biotypes, respectively [Figure 7B]. This disruption was dominated by accumulating metabolites since less than 1% of the metabolites were reduced across the biotypes. Further, within each CANOPUS superclass, the number of increased metabolites was higher for the GR-biotypes than GS-biotypes except for lipids and lipid-like molecules, where GS-biotypes showed an accumulation of 62 metabolites compared to 45 in the GR-biotypes [Figure 7B].

Within identified secondary metabolites, the GR-biotypes showed an increase in the abundance of three phenylpropanoids [Figure 8] and a terpenoid [Figure 9], while another ferulic acid derivative was reduced in the drought-treated GR-biotypes. Interestingly, there were no significant perturbations in the measured phenylpropanoid and terpenoid profile of drought-treated GS-biotypes [Figure 8, 9, Table S4]; however, we did observe a reduction in the abundance of hypoxanthine pentoside [Figure 9, Table S4].

## DISCUSSION

### Innate metabolic differences between the GS- and GR-biotypes of Palmer amaranth

Despite high EPSPS gene copy numbers, the abundance of shikimate pathway-derived metabolites, including aromatic amino acids, phenylpropanoids, and phenolic compounds, was not higher in the non-stressed GR-biotypes. Our results are in line with previous reports that observed a similar abundance of aromatic amino acids when comparing single GS- and GR- biotype (Maroli et al., 2015; Fernández-Escalada et al., 2016, 2017). This indicates that even when abundant (Gaines et al., 2010; Ribeiro et al., 2014; Fernández-Escalada et al., 2016), the catalytic activity of the EPSPS enzyme in the GR-biotypes could be limited by substrate supply or feedback inhibition (Maeda and Dudareva, 2012), or a combination thereof. Interestingly, the GS-biotypes had twice the number of phenylpropanoids (and other unidentified phytochemicals) with higher native abundance compared to GR-biotypes. Specifically, GS-biotypes could be distinguished based on the high abundance of coumaric acid amides and glycosides of kaempferol and quercetin. Variation in the abundance of hydroxycinnamic acid conjugates and flavonoids has been observed across *Amaranthus* sp as a function of biotypes, species, and growing conditions (Steffensen et al., 2011; Niveyro et al., 2013; Pedersen et al., 2010; Sarker and Oba, 2019; 2020). Overall, our results show that, despite the potential of EPSPS-mediated upregulation of the phenylpropanoid pathway, the native abundance of defense compounds produced from this pathway, under non-stressed conditions, is lower in GR-biotypes.

In contrast to shikimate pathway metabolites, GR-biotypes did show a higher abundance of non-aromatic amino acids and sugars than GS-biotypes, which suggests a higher activity of central carbon metabolism in the non-stressed GR-biotypes. This higher energy metabolism could, in turn, defray the significant energy costs involved in gene duplication, transcription, and translation of EPSPS (Vila-Aiub et al., 2019) without suffering any pleiotropic effect of the EPSPS amplification (Giacomini et al., 2014; Vila-Aiub et al., 2014).

### Differential effect of glyphosate on GS- and GR-biotypes of Palmer amaranth

Exposure to glyphosate induced phytochemical alterations across GS- and GR-biotypes, but the two groups differed with respect to the extent of disruption. Accumulation of shikimic acid was observed in both groups – higher in GS-biotypes – confirming previous reports (Nandula et al., 2012; Maroli et al., 2015; Fernández-Escalada et al., 2019). The initial disruption of the shikimate pathway in GR-biotypes points to a blockage of the EPSPS activity, despite the possibility of an additional pool of the EPSPS enzyme (through EPSPS amplification) (Gaines et al., 2010; 2011). Previous studies using a single GR-biotype have documented the shikimate accumulation at 8HAT and subsequent recovery of plants by 80HAT (Maroli et al., 2015). The present study confirms that the glyphosate-induced initial blockage of EPSPS is common to other GR-biotypes and points to the possibility that the excess amount of EPSPS might be functionalized only after the herbicide application, and the associated time lag in the enzyme induction results in the transient accumulation of shikimic acid. Disruption of central carbon metabolism (amino acids, TCA cycle metabolites, and sugars) also provides evidence towards glyphosate-induced stress in GR-biotypes (Maroli et al., 2015; Fernández-Escalada et al., 2016; 2019). However, these perturbations of primary metabolism were significantly lower than those observed in GS-biotypes, indicating that GR-biotypes are not entirely unprotected from glyphosate.

Further, more than 80% of the glyphosate-induced increase in the abundance of phenylpropanoids (and other unidentified phytochemicals) in GS-biotypes was also observed in GR-biotypes, revealing that the glyphosate-induced chemotype of GS-biotypes is dominated by specificity to herbicide stress. However, GR-biotypes did exhibit a significant group-specific phytochemical response to glyphosate. Specifically, the glyphosate-induced accumulation of phenylpropanoids was higher in GR-biotypes than GS-biotypes holding our second hypothesis only partially true. Phenylpropanoids form an essential component of the phytochemical protection machinery of plants owing to their antioxidant and reactive oxygen species (ROS) scavenging abilities (Deng and Lu, 2017; Sharma et al., 2019). Thus, the higher accumulation of these compounds in GR-biotypes following glyphosate application could contribute to the protection and recovery of glyphosate-treated GR-biotypes until the activation of the excess EPSPS enzyme. This induced antioxidant capacity can also help combat the oxidative stress induced by general stressors (Mittler, 2002; Böttger et al., 2018) that could prime the GR- biotypes against non-optimal growing conditions post-exposure to the non-lethal herbicide dose.

In addition to increased abundance, the composition of phenylpropanoids in GR-biotypes was also modulated towards more efficient antioxidants. Specifically, the glyphosate-treated GR- biotypes showed an increased abundance of glycosides of dihydroxylated (quercetin) and methoxylated flavanols (isorhamnetin), compared to monohydroxylated flavanol, kaempferol in the GS-biotypes. The dihydroxylated flavanols are known to exhibit stronger antioxidant capacity than monohydroxylated flavanols (Mierziak et al., 2014; Alseekh et al., 2020), while methoxylation provides higher membrane permeability and stability to isorhamnetin (Wen and Walle, 2006; Alseekh et al., 2020).

Further, despite the specific inhibition of EPSPS enzyme by glyphosate, the impact of glyphosate extended beyond the shikimate (and phenylpropanoid pathway), as displayed by the accumulation of terpenoids across the groups, rejecting our third hypothesis. This could be the result of the secondary or off-target effects of glyphosate. However, in contrast to phenylpropanoids, the accumulation of triterpenoid glycosides was similar across the groups indicating that glyphosate-induced phytochemical differences between the GS- and GR-biotypes are limited to the pathways downstream of the herbicide target. Thus, the glyphosate-induced phytochemical response of conspecifics with the differential capability to tolerate glyphosate had both shared and specific components where the group-specific response is dominated by metabolites produced through the targeted shikimate pathway.

### Differential effect of drought treatment on GS- and GR-biotypes of Palmer amaranth

Droughted GS- and GR-biotypes of Palmer amaranth sustained only minor metabolic disruptions, a pattern divergent from glyphosate treatment. This indicates that the identity of the stressor was a driving factor in producing stress-induced phytochemical changes. The impact of drought was minimal on TCA cycle metabolites across the groups, whereas amino acids were disrupted only in GR-biotypes and sugars only in GS-biotypes. Similar to primary metabolism, the impact of drought treatment on the phenylpropanoid and terpenoid pathways was also negligible. This indicates that the EPSPS amplification-mediated activation of the phenylpropanoid pathway in GR-biotypes did not occur after perception of a general stressor drought; instead, the accumulation of phenylpropanoids was a specific response to a non-lethal dose of glyphosate. Further, the phytochemical perturbations induced by drought were higher in GR-biotypes than in GS-biotypes. This, along with the innate differences between the GS- and GR- biotypes, suggest that the cost of EPSPS amplification in Palmer amaranth could be imposed in terms of reduced investment in defense compounds, and thus, the fitness cost of EPSPS needs to be evaluated under sub-optimal growing conditions. Kohrt et al. (2016) also reported a potential effect of fertilization and soil moisture status on the glyphosate sensitivity of GR plants under field conditions.

Taken together, our study provides strong evidence towards the inducibility and specificity of glyphosate-induced phytochemical response in GR Palmer amaranth. The inclusion of multiple biotypes as a random factor within GS- and GR- groups, by accounting for differential genetic background of biotypes, shows that the partial inducibility of glyphosate resistance is a common phenomenon across populations of Palmer amaranth. Further, by initiating the accumulation of phytochemicals, the non-lethal doses of glyphosate could act as priming agents in the GR-biotypes, which could enhance the ability of GR-biotypes to tolerate future stress events. Overall, it can be concluded that the identity of stressor dominates the glyphosate stress tolerance capabilities across biotypes of Palmer amaranth in inducing a signature chemotype, i.e., the phytochemical response is stress-specific rather than group-specific.

## EXPERIMENTAL PROCEDURES

### Plant material and growth conditions

Three GR (CoR, C1B1, and T4B1) and two GS (CoS and S17) biotypes of Palmer amaranth were used in this study. Palmer amaranth was selected due to its economic importance in row crop production systems (*Zea mays*, *Glycine max*, and *Gossypium hirsutum*) of the southeastern United States (US) where it causes yield losses ranging from 50-90% (Ward et al., 2013; Lindsay et al., 2017). The CoR biotype with GR_50_ of 1.6 kg ha^-1^ and EPSPS gene copy number (relative to ALS) of >30 (for >90% of the population) (Gaines et al., 2010; 2011) and the CoS biotype (GR_50_: 0.04 kg ha^-1^; EPSPS gene copy number: 1 (Gaines et al., 2010; 2011)) were collected in Georgia, US (provided by Todd Gaines, Colorado State University, Fort Collins, CO) while C1B1 (GR_50_: 1.5 kg ha^-1^, EPSPS gene copy number >55 (Nandula et al., 2012; Ribeiro, 2014)), T4B1 (GR_50_: 1.3 kg ha^-1^, EPSPS gene copy number >33 (Nandula et al., 2012; Ribeiro, 2014)), and S17 (GR_50_: 0.09 kg ha^-1^, EPSPS gene copy number: 1 (Nandula et al., 2012; Ribeiro, 2014)) were collected in Mississippi, US (USDA-ARS, Stoneville, MS). The seeds were sown in 48-cell trays in the germination mix (SunGro). After one week, the seedlings were transplanted to pots (10 cm diameter x 9 cm deep) containing germination mix. Plants were watered every other day before the drought application and kept in the greenhouse for the duration of the experiment. The pots were fertilized four days after transplanting with 50ml of 4g L^-1^ fertilizer (MiracleGro, 24%−8%−16% [N−P−K], Scotts Miracle-Gro Products, Inc., Marysville, OH).

### Treatment application

Twelve days after transplanting, the drought treatment was initiated in one-third of the plants by withholding the irrigation. Drought treatment was maintained at 22 ± 8% water holding capacity (WHC), while the non-drought treatments were kept at 80 ± 5% WHC by monitoring the water content of the pots and irrigating them daily. Three days after the initiation of drought treatment, a separate one-third of regularly watered plants were treated with glyphosate. In glyphosate treatment, plants were sprayed with half the field recommended dose of glyphosate (0.42 kg ae ha^-1^ Roundup ProMax, Bayer CropScience, St. Louis, MO) using a Teejet 8001EVS spray nozzle in an enclosed spray chamber (DeVries Manufacturing, Hollandale, MN). The remaining one-third of the plants that received neither the drought treatment nor the glyphosate application served as a control in each biotype. Samples were harvested 44 hours after treatment with glyphosate or five days after the initiation of drought treatment. Three hours before harvest, all the plants were watered to bring the moisture content within the respective treatment level, and five biological replicates from each treatment were randomly selected for sampling. The young, fully opened meristematic leaves were collected into weighed tubes and stored at -80°C until analysis. At the time of harvest (44 hours after glyphosate application), the glyphosate treated CoR, C1B1, T4B1, and S17 did not show any visual toxicity symptoms, while the CoS exhibited slight leaf curling.

### Metabolite extraction

Metabolites were extracted from leaves to analyze the primary and secondary metabolite profile of samples using GC-MS and UHPLC-MS/MS, respectively, according to Narvekar and Tharayil (2021). To ∼100 mg weighed leaf tissue, methanol was added in a ratio of 1:10 (w/v), followed by homogenization at 6000 rpm for 30 seconds for four cycles at 2°C. Samples were centrifuged at 2500 rpm for 4 minutes, and 500 µL of supernatant was transferred to a new 2 mL tube. Ice-cold chloroform was added to the 500 µL sample extract, followed by cold water at a ratio of 1:1:1 (methanol extract/chloroform/water, v/v/v) and kept in ice for 10 minutes for phase separation. After centrifugation at 10000 rpm for one minute, top methanol-water was used for instrument analysis.

### Untargeted primary metabolomics analysis by GC-MS

Chloroform partitioned methanol-water extract from above was used to profile polar primary metabolites using GC-MS, according to Maroli et al. (2015). Briefly, 50 µL of the methanol-water extract was transferred into vials with glass inserts, 10 µL of 50 μg mL^−1^ ribitol (internal standard), and 20 μg mL^−1^ of d27-myristic acid (retention time lock) in hexane were added to the vials and dried down under a nitrogen stream. The dried samples were methoxylaminated at 40°C with 20 μL of methoxylamine hydrochloride (20 mg mL^−1^) in pyridine for 90 minutes followed by silylation of the metabolites with 90 μL of N-methyl-N-(trimethylsilyl) trifluoroacetamide (MSTFA) with 1% trimethylchlorosilane (TMCS) for 30 minutes at 40°C. The derivatized metabolites were separated by gas chromatography (Agilent 7980, Agilent Technologies, Santa Clara, CA) on a J&W DB-5 ms column (30 m × 0.25 mm × 0.25 μm, Agilent Technologies) and analyzed using a transmission quadrupole mass spectrometer (Agilent 5975 C series) with an electron ionization interface. An injection volume of 1 µL was used, and the carrier gas, helium, was operated in constant pressure mode at 7.66 psi. The GC inlet and auxiliary temperatures were set to 270°C and 285°C, respectively, with a split ratio of 5:1. The oven temperature gradient began with a hold at 60°C for one minute, followed by a 10°C/min ramp to 300°C, and hold at 300°C for five minutes. Ions were produced by EI at 70 eV and analyzed in quadrupole with a scan range of m/z 60-460 with a detector gain factor of one. Pooled samples were run as quality control (QC). Mass spectra were processed on MS-Dial (Tsugawa et al., 2011a; 2011b; Matsuo et al., 2017). The identification of the primary metabolites was based on the Kovats RI, and the mass fragmentation pattern matches with Kovats RI Library (GCMS DB_AllPublic-KovatsRI-VS2) with a minimum EI similarity score of 80%.

### Untargeted secondary metabolomics analysis by UHPLC-MS/MS

#### a. Instrument analysis

150 µL of the top methanol-water phase was transferred into a polypropylene vial insert containing 150 µL of the internal standard, ^13^C_6_ resveratrol (0.5 ng/µL), and was analyzed on UHPLC-MS/MS for capturing the secondary metabolite profile. Pooled samples and method blanks were run as QC. Metabolites were separated on an Acclaim PepMap 100 C18 (150 x 1 mm, 3 μm) column maintained at 32°C, using an Ultimate 3000 UHPLC (ultra-high pressure liquid chromatography) coupled to an Orbitrap Fusion Tribrid mass spectrometer equipped with heated electrospray ionization (HESI-II) source (Thermo Scientific, Waltham, MA, USA). Mobile phases A and B consisted of 0.05% formic acid in water for A and in acetonitrile for B. The solvent gradient started with a two-minute hold at 5% B with a flow rate of 0.07 mL/min, increased to 95% B at 14.5 minutes with a flow rate of 0.12 mL/min, and returned to 5% B for four minutes with a flow rate of 0.07 mL/min. The HESI ion source was operated in positive ionization mode, with spray voltage set to 3.5 kV, ion transfer tube and vaporizer temperatures of 300°C and 150°C, respectively, and sheath, auxiliary, and sweep gases set to 1.6 mL/min, 12 mL/min, and 0.75 mL/min, respectively. The injection volume was 4 µL.

For quantitation and identification of metabolites in the samples, data-dependent MS^2^ was collected using Acquire X background exclusion method (Xcalibur software, Thermo Scientific). MS^1^ scan data was collected in orbitrap at 60 K resolution with 2e5 automated gain control (AGC) and 50 ms maximum injection time for precursor ions within the scan range of m/z 120-1000. Precursor ions that passed the filters (intensity threshold of 1e5, dynamic exclusion duration of 4 seconds for 10 ppm mass tolerance, and exclusion mass list automatically generated by the AcquireX background exclusion workflow) were sent to orbitrap for MS^2^ acquisition. For MS^2^ fragmentation, stepped high energy C-trap dissociation (HCD) of 20, 35, 50% was used, and the scan data was collected in orbitrap at 7.5 K resolution with 1e5 AGC and 22 ms maximum injection time. The pooled sample was also analyzed in negative ionization mode (spray voltage of 2.6 kV) with data-dependent MS^2^ for compound identification.

#### b. Compound identification

For quantitation and identification of compounds, mass spectra obtained from UHPLC-MS/MS were processed in a node-based workflow of Compound Discoverer v3.1 (Thermo Fisher Scientific, MA, USA). The compounds were annotated based on accurate mass match (error < 5 ppm) and characteristic fragmentation patterns of compounds in various online libraries (MoNA, mzCloud, Flavanoid Search, HMDB) and literature. Compound IDs were also confirmed based on the compound class prediction of CANOPUS and CSI:Finger ID from the SIRIUS v4.4 analysis (Dührkop et al., 2019). Additionally, the fragmentation pattern of triterpenoid glycosides was confirmed by running two reference standards, Ginsenoside Ro (CAS No. 34367-04-9) and Momordin Ic (CAS No. 96990-18-0), under the same UHPLC-MS/MS conditions. The mass spectra (MS^1^ and MS^2^) of standards are given in supplementary figures [Figure S6; S7].

#### c. CANOPUS analysis

The mass spectrometric peaks obtained from UHPLC-MS/MS analysis were also classified using CANOPUS (Dührkop et al., 2021). CANOPUS is a computational tool built within SIRIUS v4.4 (Dührkop et al., 2019) that predicts the classification of compounds based on fragmentation patterns into a hierarchical classification of ClassyFire (Djoumbou Feunang et al., 2016). For processing, the mzML files for each sample were imported into the SIRIUS v4.4. The peaks eluting before 2.00 minutes and after 14.50 minutes were removed prior to analysis. The features were scored for MS/MS isotopes (Böcker et al., 2009) with an MS^2^ mass deviation of 5 ppm. The elemental composition was limited to molecular formulas available in databases. [M+Na]+, [M+K]+, and [M+H]+ adducts were allowed for molecular formula prediction while [M+NH4]+ and [M+CAN+H]+ were additionally included as fallback adducts for CSI:Finger ID (Dührkop et al., 2015; Böcker and Dührkop, 2016; Hoffmann et al., 2021). ZODIAC (Ludwig et al., 2020) was also used to improve the annotation of molecular formulas.

### Statistical analysis

Peak areas were used as a measure of metabolite abundance. Due to the high degree of insource fragmentation observed in triterpenoid glycosides (Ma et al., 2019) [Figure S6], the highest intensity peak was used for the quantitation. We observed linearity in the peak areas of the quantified insource fragment and parent compound [Figure S8]. Before statistical analysis, biotypes within each group (GS and GR) were grouped together. The multivariate analysis was carried out in Metaboanalyst v5.0 (Pang et al., 2021). The data was autoscaled before analysis on Metaboanalyst to meet the assumption of normality. The statistical significance of metabolites was determined by performing univariate analysis (parametric t-test) between the control and treatments (drought/glyphosate) with a significance level of *p*<0.05 in the GraphPad Prism v9.0 (GraphPad Software, La Jolla, CA). Errors due to multiple comparisons were controlled by maintaining the false discovery rate (FDR) <0.05 using the two-stage step-up method of Benjamini, Krieger, and Yekutieli (Benjamini et al., 2006). The graphs were plotted using GraphPad Prism v9.0 and Microsoft Excel 2016 (Microsoft Corporation, One Microsoft Way, Redmond, WA).

## SUPPLEMENTARY INFORMATION

**Figure S1:** VIP score plots for the first two components from sPLS-DA of identified primary metabolites in GS- and GR-biotypes.

**Figure S2:** Correlation plot representing correlation coefficients of the top 25 identified primary metabolites with shikimic acid.

**Figure S3:** sPLS-DA and VIP score plot of GS- and GR- biotypes in control treatment for identified primary metabolites.

**Figure S4:** Native differences in the abundance of identified primary metabolites between GS- and GR-biotypes.

**Figure S5:** Effect of glyphosate and drought treatment on total sugar metabolites in GS- and GR-biotypes.

**Figure S6:** MS^1^ spectrum of standard and identified triterpenoids.

**Figure S7:** MS^2^ spectrum of standard and identified triterpenoids.

**Figure S8:** Linearity in the peak areas for the highest abundant insource fragment and parent m/z for triterpenoids.

**Table S1:** Primary metabolites identified in Palmer amaranth from GC-MS analysis.

**Table S2:** Pairwise comparisons of identified primary metabolites between control, glyphosate-, and drought-treated GS- and GR-biotypes of Palmer amaranth.

**Table S3:** Secondary metabolites identified in Palmer amaranth from UHPLC-MS/MS analysis. **Table S4:** Pairwise comparisons of identified secondary metabolites between control, glyphosate-, and drought-treated GS- and GR-biotypes of Palmer amaranth.

**Table S5:** CANOPUS classification results and pairwise comparisons of filtered featured from UHPLC-MS/MS analysis.

