## Supplementary Figures for "Metabolite profiles across populations of Palmer amaranth (*Amaranthus palmeri*) highlight the specificity and inducibility of phytochemical response to glyphosate stress"

**Figure S1: VIP score plots for first two components from sPLS-DA of identified primary metabolites in GS- and GR-biotypes**: Variable importance in Projection (VIP) score plot for the (A) first and (B) second principal component from the sPLS-DA plot of identified primary metabolites in glyphosate- (Gly), drought- (Drt) and control- (Amb) treated GR- and GS- biotypes of Palmer amaranth. Within each group (GR & GS) the biotype identity is a random factor.

**Figure S2: Correlation plot representing correlation coefficients of the top 25 identified primary metabolites with shikimic acid.**

**Figure S3: sPLS-DA and VIP score plot of GS- and GR-biotypes in control treatment for identified primary metabolites:** sPLS-DA clustering (left) and VIP score plot for first component (right) of non-treated glyphosate-susceptible (GS) and -resistant (GR) biotypes of Palmer amaranth. Within each group (GR & GS) the biotype identity is a random factor.

**Figure S4: Native differences in the abundance of identified primary metabolites between GS- and GR-biotypes.** $Log_2$ fold change plot of identified primary metabolites that were significantly different ($Log_2$ fold change > |1|, $p < 0.05$, FDR <0.05) between the control GR (GR_Amb)- and GS (GS_Amb)-biotypes. GR_Amb and GS_Amb are represented by green and yellow color respectively. Values in each cell are the GR_Amb/GS_Amb $log_2$ fold change values. Within each group (GS & GR) the biotype identity is a random factor.

**Figure S5: Effect of glyphosate and drought treatment on total sugar metabolites in GS- and GR-biotypes.** $Log_2$ fold change in total sugar metabolites (glucose, ribose, arabinose, fructose, galactose, sucrose, tagatose, myo-inositol and galactinol) in glyphosate- (Gly) and drought- (Drt) treated GR- and GS- biotypes of Palmer amaranth. Bar represents mean + SD. Asterisk denotes the significant difference of treatment compared to control at $p < 0.05$ (Tukey's HSD). Bars with different letters are significantly different from each other at $p < 0.05$ (Tukey's HSD). Within each group (GR & GS) the biotype identity is a random factor.

**Figure S6: $MS^1$ spectrum of standard and identified triterpenoids.** $MS^1$ spectrum of $[M+H]^+$ of standard and identified terpenoids showing insource fragmentation for triterpenoid standards A) Ginsenoside Ro B) Momordin lc and two identified triterpenoid glycosides: Oleanolic acid-3-pentopyranoosylhexopyranosylhexopyrauronoside (C) and Medicagenic acid-3-pentopyranosylhexopyranosylhexopyrauronoside (D) observed in Palmer amaranth.

**Figure S7: $MS^2$ spectrum of standard and identified triterpenoids.** $MS^2$ spectrum of $[M+H]^+$ for triterpenoid standards A) Ginsenoside Ro B) Momordin lc and two identified triterpenoid glycosides: Oleanolic acid-3-pentopyranoosylhexopyranosylhexopyrauronoside (C) and Medicagenic acid-3-pentopyranosylhexopyranosylhexopyrauronoside (D) observed in Palmer amaranth. Inset shows the fragmentation pathway.

**Figure S8: Linearity in the peak areas for the highest abundant insource fragment and parent m/z for triterpenoids.** A) Oleanolic acid-3-pentopyranoosylhexopyranosylhexopyrauronoside (B) Medicagenic acid-3-pentopyranosylhexopyranosylhexopyrauronoside observed in Palmer amaranth.

**Figure S1: VIP score plots for first two components from sPLS-DA of identified primary metabolites in GS- and GR-biotypes**: Variable importance in Projection (VIP) score plot for the (A) first and (B) second principal component from the sPLS-DA plot of identified primary metabolites in glyphosate- (Gly), drought- (Drt) and control- (Amb) treated GR- and GS-biotypes of Palmer amaranth. Within each group (GR & GS) the biotype identity is a random factor.

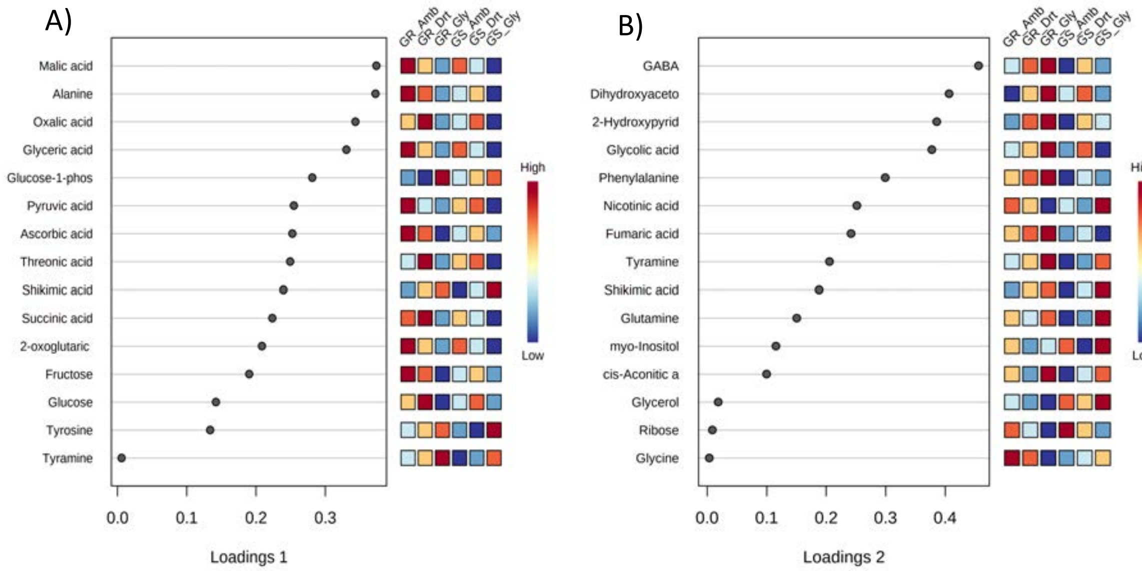


**Figure S2: Correlation plot representing correlation coefficients of the top 25 identified primary metabolites with shikimic acid.**

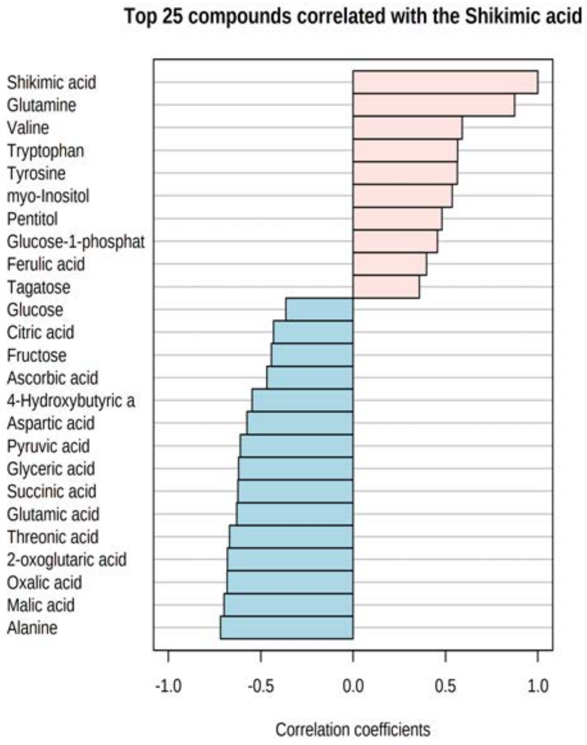

**Figure S3: sPLS-DA and VIP score plot of GS- and GR-biotypes in control treatment for identified primary metabolites:** sPLS-DA clustering (left) and VIP score plot for first component (right) of non-treated glyphosate-susceptible (GS) and -resistant (GR) biotypes of Palmer amaranth. Within each group (GR & GS) the biotype identity is a random factor.

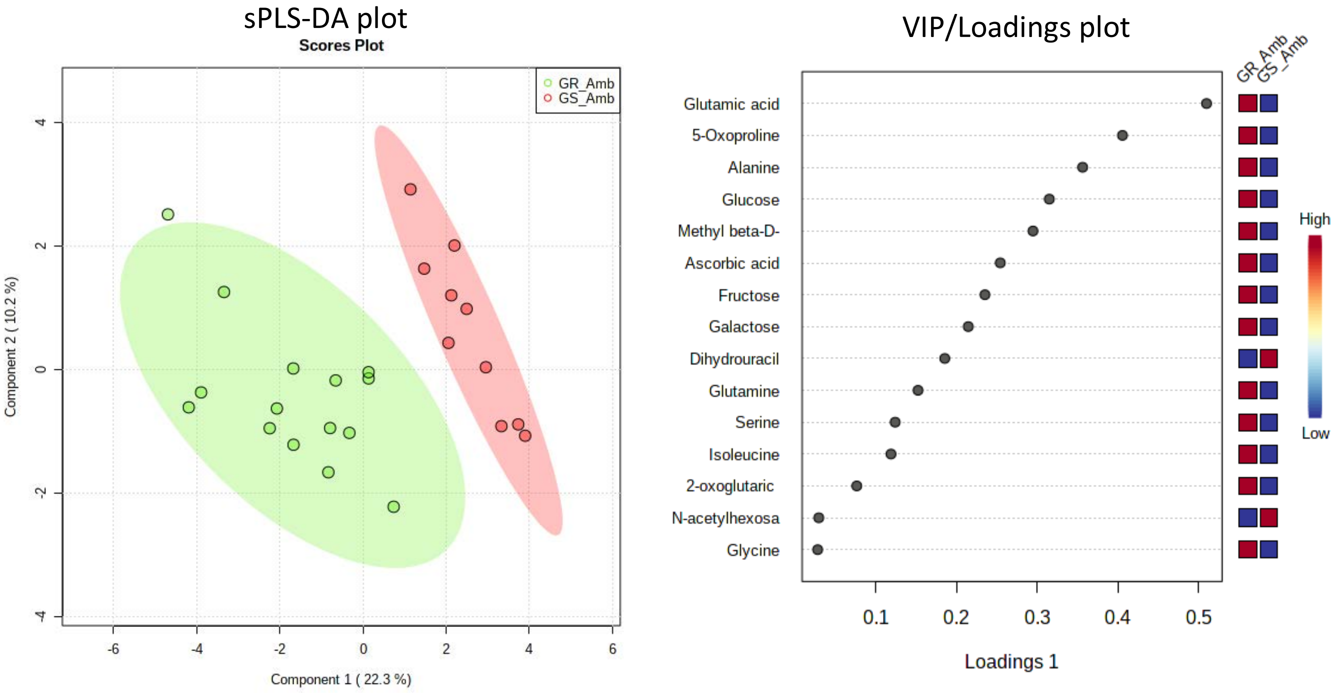

**Figure S4: Native differences in the abundance of identified primary metabolites between GS- and GR-biotypes.** $Log_2$ fold change plot of identified primary metabolites that were significantly different ($Log_2$ fold change > |1|, p <0.05, FDR <0.05) between the control GR (GR_Amb)- and GS (GS_Amb)-biotypes. GR_Amb and GS_Amb are represented by green and yellow color respectively. Values in each cell are the GR_Amb/GS_Amb $log_2$ fold change values. Within each group (GS & GR) the biotype identity is a random factor.

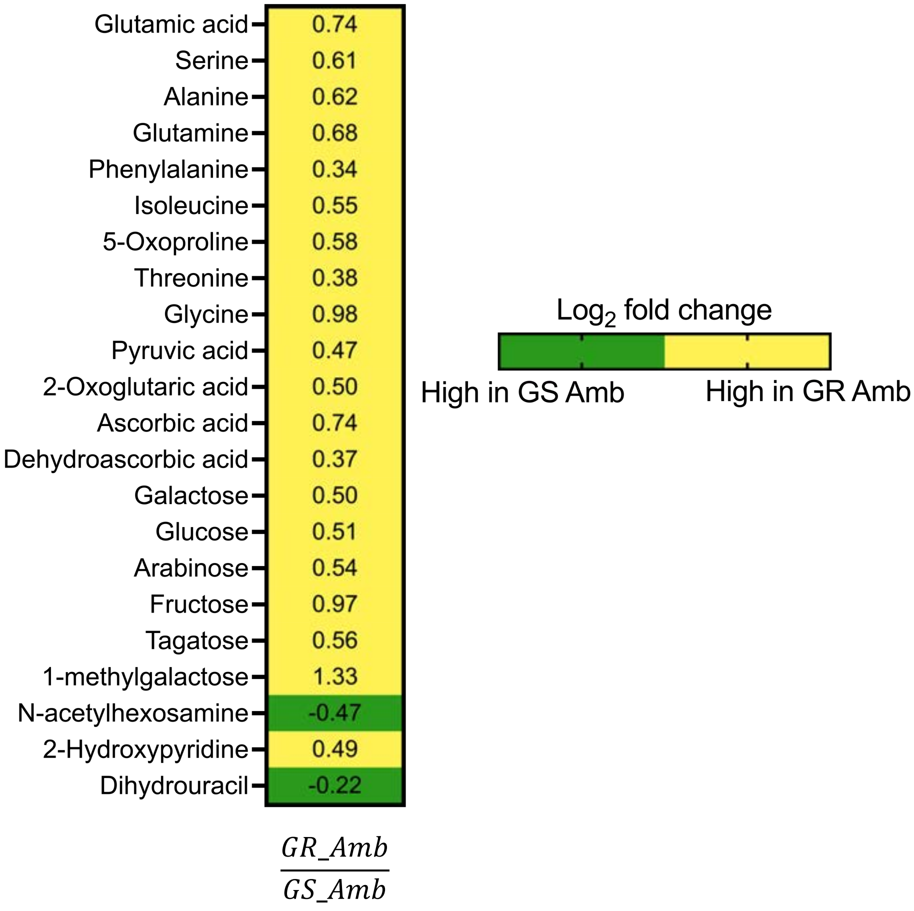

**Figure S5: Effect of glyphosate and drought treatment on total sugar metabolites in GS- and GR-biotypes.** $Log_2$ fold change in total sugar metabolites (glucose, ribose, arabinose, fructose, galactose, sucrose, tagatose, myo-inositol and galactinol) in glyphosate- (Gly) and drought- (Drt) treated GR- and GS- biotypes of Palmer amaranth. Bar represents mean + SD. Asterisk denotes the significant difference of treatment compared to control at $p < 0.05$ (Tukey's HSD). Bars with different letters are significantly different from each other at $p <0.05$ (Tukey's HSD). Within each group (GR & GS) the biotype identity is a random factor.

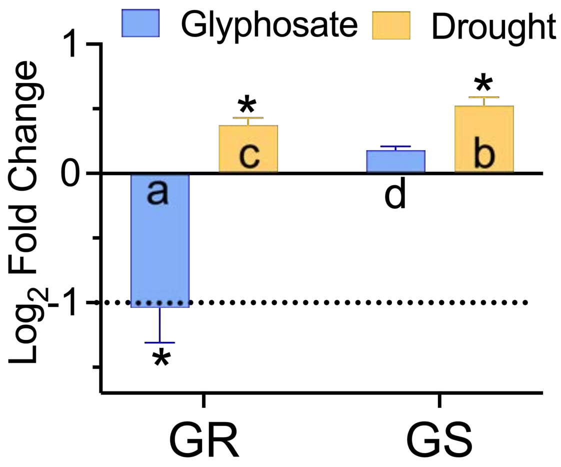

**Figure S6: MS[1] spectrum of standard and identified triterpenoids.** MS[1] spectrum of $[M+H]^+$ of standard and identified terpenoids showing insource fragmentation for triterpenoid standards A) Ginsenoside Ro B) Momordin Ic and two identified triterpenoid glycosides: Oleanolic acid-3-pentopyranoosylhexopyranosylhexopyrauronoside (C) and Medicagenic acid-3-pentopyranosylhexopyranosylhexopyrauronoside (D) observed in Palmer amaranth.

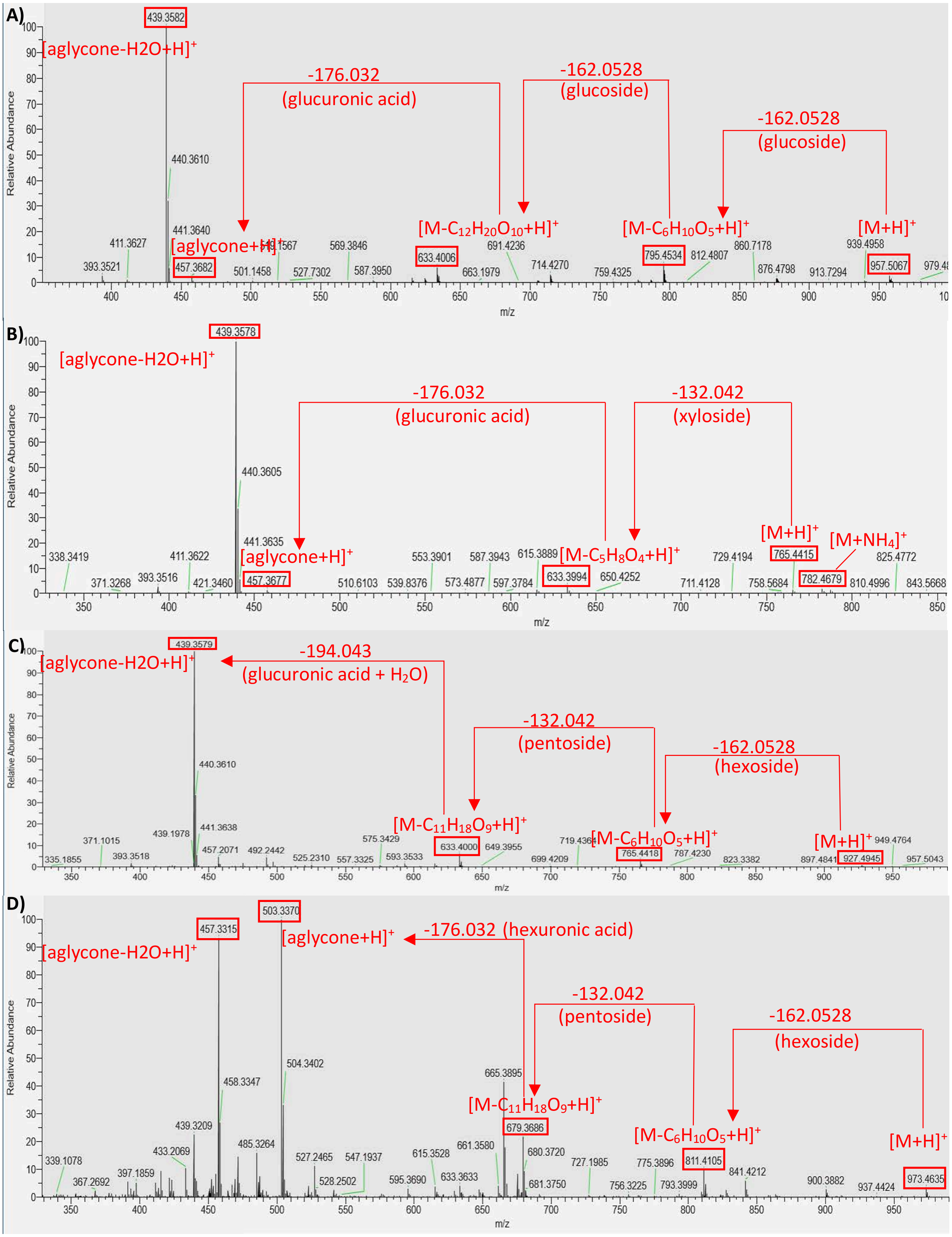

**Figure S7: $MS^2$ spectrum of standard and identified triterpenoids.** $MS^2$ spectrum of $[M+H]^+$ for triterpenoid standards A) Ginsenoside Ro B) Momordin lc and two identified triterpenoid glycosides: Oleanolic acid-3-pentopyranoosylhexopyranosylhexopyrauronoside (C) and Medicagenic acid-3-pentopyranosylhexopyranosylhexopyrauronoside (D) observed in Palmer amaranth. Inset shows the fragmentation pathway.

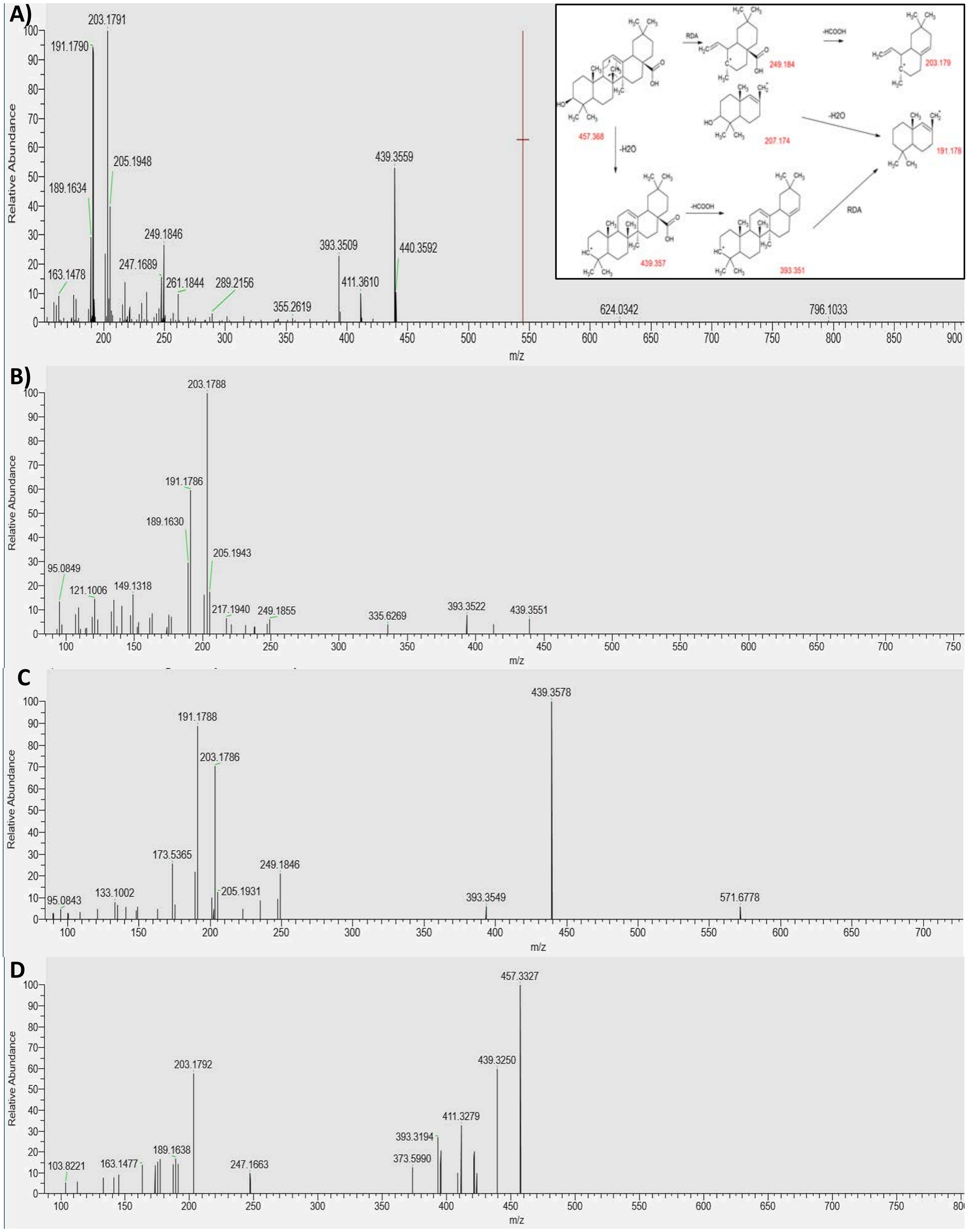

**Figure S8: Linearity in the peak areas for the highest abundant insource fragment and parent m/z for triterpenoids.** A) Oleanolic acid-3-pentopyranoosylhexopyranosylhexopyrauronoside (B) Medicagenic acid-3-pentopyranosylhexopyranosylhexopyrauronoside observed in Palmer amaranth.

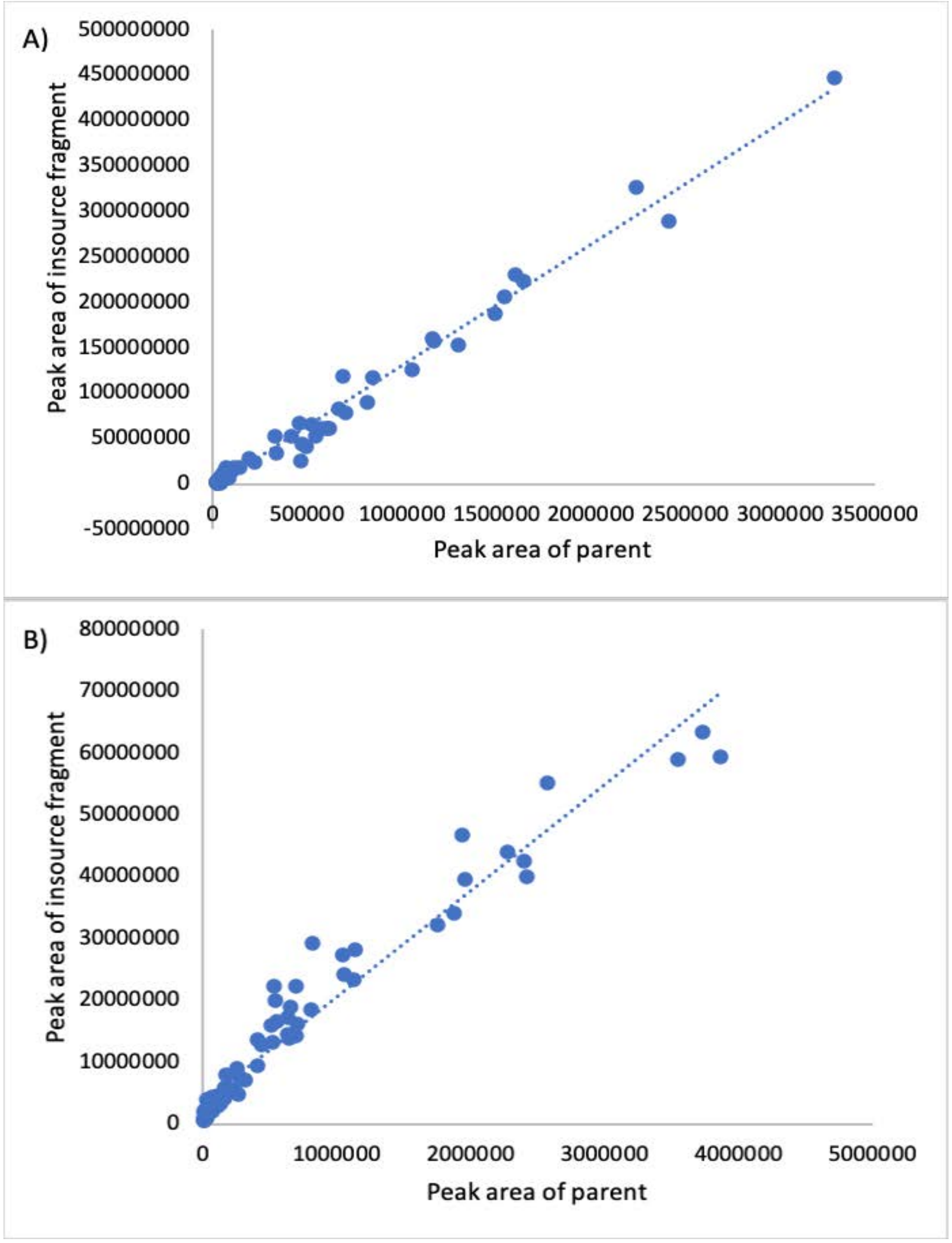
